# A Dual Role for LAR-RPTP in Regulating Long-distance Transport and Synaptic Retention of AMPARs, Essential for Long Term Associative Memory

**DOI:** 10.1101/2022.08.11.503694

**Authors:** D.M. Pierce, Z. Lenninger, R.L. Doser, K.M. Knight, A. Stetak, F.J. Hoerndli

**Affiliations:** Sacramento City College; Colorado State University; University of Basel, Biozentrum

## Abstract

The AMPA subtype of ionotropic glutamate receptors (AMPARs) plays an essential role in excitatory synaptic transmission, learning, and memory. The majority of AMPARs are made in the cell body and are transported by molecular motors to synapses. Maintaining the proper number of synaptic receptors requires coordinated regulation of receptor production, export from the soma and delivery at synapses. This major logistical process is essential for circuit function and behavior. Although recent studies have shown that long-distance synaptic transport is regulated by neuronal activity, little is known about the mechanisms that coordinate somatic export or synaptic delivery and removal. Here we show that loss of the PTP-3A isoform of the receptor tyrosine phosphatase PTP-3 (the *C. elegans* homologue of vertebrate LAR-RPTP) leads to a ∼60% decrease in AMPAR transport; this affects synaptic delivery of AMPARs and synaptic functions necessary for long-term associative olfactory memory in *C. elegans*. Interestingly, while complete loss of PTP-3A leads to defects in transport and local synaptic trafficking of AMPARs, loss of only PTP-3 phosphatase function affects local synaptic recycling and retention of AMPARs. Finally, we show that the N-terminus of PTP-3A regulates transport, whereas the C-terminal regulates synaptic retention of AMPARs. Altogether, our results suggest a model in which the two domains of PTP-3/LAR-RPTPs have specific, complementary roles in coordinating somatic export and local retention of AMPARs essential for long-term associative memory.

## INTRODUCTION

The AMPA subtype of ionotropic glutamate receptors (AMPARs) is the workhorse of excitatory synaptic transmission. Regulating the quantity of AMPARs at synapses is essential for both input-dependent synaptic plasticity and network-dependent homeostatic plasticity (Diering and Huganir, 2018; Groc and Choquet, 2020; Grochowska et al., 2022). However, the majority of AMPARs are produced in the cell body and therefore must be transported to the synapses (Hanus and Ehlers, 2016). This complex, multistep process is regulated on many levels to finely tune the form and strength of synapses. Long-distance transport of AMPARs by molecular motors is one essential process in the distribution of receptors. However, this is the least understood step in AMPAR trafficking. New studies have started to reveal that AMPAR transport is regulated by neuronal activity, namely increases in cytoplasmic calcium and calcium/calmodulin-dependent protein kinase II (CaMKII) (Doser et al., 2020; Hangen et al., 2018; Hoerndli et al., 2015). Although many mechanistic details of how this pathway exerts control over transport remain unidentified, these studies have revealed that phosphorylation of both the cargo (i.e., AMPARs) and motor-adaptors contribute to long-distance transport regulation (Hangen et al., 2018; Hoerndli et al., 2015).

Calcium signaling and CaMKII activation play essential roles in transport (Hangen et al., 2018; Hoerndli et al., 2015). Although calcium signaling is certainly a central regulator of synaptic AMPAR numbers, other signaling pathways are known. For instance, tyrosine phosphorylation has been recently identified as a regulator of synaptic and homeostatic plasticity (Gladding et al., 2009; Lua et al., 2010; Han L. Tan et al., 2020; Yong et al., 2020). Phosphorylation and dephosphorylation of a single tyrosine residue in the AMPAR GluA2 subunit were observed to respectively up or downscale homeostatic plasticity *in vivo* (Yong et al., 2020) and has been shown to be critical for AMPAR recruitment in long-term potentiation (LTP) (Tan et al., 2020). Interestingly, tyrosine phosphatases, such as Receptor Tyrosine Phosphatases of the leukocyte common antigen-related family (LAR-RPTPs), have been documented to recruit and stabilize AMPARs at synapses during development (Dunah et al., 2005), but the exact mechanism of how that is achieved is not understood. One possibility is that LAR-RPTPs can bind to Glutamate Receptor Interacting Protein 1 (GRIP-1) forming a complex with the scaffold Liprin-α (Shin et al., 2003). GRIP-1 and Liprin-α have been shown to bind to the molecular motor, Kinesin-1, and it has been suggested that CaMKII phosphorylation of Liprin-α releases LAR-RPTP from the complex to be recruited to synapses (Miller et al., 2005; Setou et al., 2002; Shin et al., 2003; Wyszynski et al., 2002). However, there is no direct evidence for the involvement of LAR-RPTPs or Liprin-α in regulating synaptic dynamics of AMPARs or transport. Therefore, new approaches to studying AMPAR transport are required to shed more light on the role of these interactions and the importance of tyrosine phosphorylation in AMPAR transport, delivery, and retention.

The type IIa subset of mammalian LAR-RPTPs includes LAR, PTP-δ and PTP-σ which are single transmembrane domain phosphatases with a large N-terminal extracellular domain containing three immunoglobulin (Ig)-like domains and a variable number of fibronectin III repeats (Coles et al., 2015; Won and Kim, 2018). The intracellular domain contains two tandem phosphatase domains (D1 and D2), with only D1 being catalytically active but D2 being important for substrate specificity (Streuli et al., 1990). LAR-RPTPs are normally cleaved post-translationally into two non-covalently linked subunits: the external E-subunit containing the Ig-like domains and fibronectin repeats and the intracellular P-subunit containing the transmembrane domain and the tandem phosphatases. In addition, variation in mRNA splicing gives rise to many different isoforms of LAR-RPTPs that have different functional effects (Han et al., 2019). Altogether, an overwhelming number of studies present a presynaptic role for LAR-RPTPs in synaptogenesis and synaptic function that are necessary for normal behavioral phenotypes (Kolkman et al., 2004; Uetani, 2000). It is thought that different class IIa LAR-RPTPs complement each other, but the lack of a viable triple genetic knock-out of all class IIa RPTPs has prevented testing of this hypothesis. However, two recent studies showed that conditional knockouts of all three class IIa LAR-RPTPs does not lead to defects in synaptogenesis, nor does it have any major effect on glutamatergic or GABAergic transmission in the hippocampus (Emperador-Melero et al., 2021; Sclip and Südhof, 2020). These recent studies further underscore our lack of understanding of the mechanisms by which LAR-RPTPs act on synaptic function.

In *C. elegans, ptp-3* is the sole type IIa LAR-RPTP gene and encodes three isoforms that differ in the extracellular domain (Figure 1A) (Harrington et al., 2002). The PTP-3A isoform is most homologous to vertebrate LAR-RPTP with three Ig-like domains and nine fibronectin repeats followed by a transmembrane domain and the tandem phosphatase domain (Figure 1A). PTP-3A is thought to be synaptic, whereas PTP-3B was found to be important in axon pathfinding and neuronal development (Ackley et al., 2005; Harrington et al., 2002). PTP-3C function has not yet been characterized. While PTP-3A has been shown to have presynaptic roles in synaptogenesis (Ackley et al., 2005; Caylor et al., 2013; Dai et al., 2006; Hartin et al., 2015), its role at postsynaptic sites in *C. elegans* has not been assessed.

**Figure 1:**
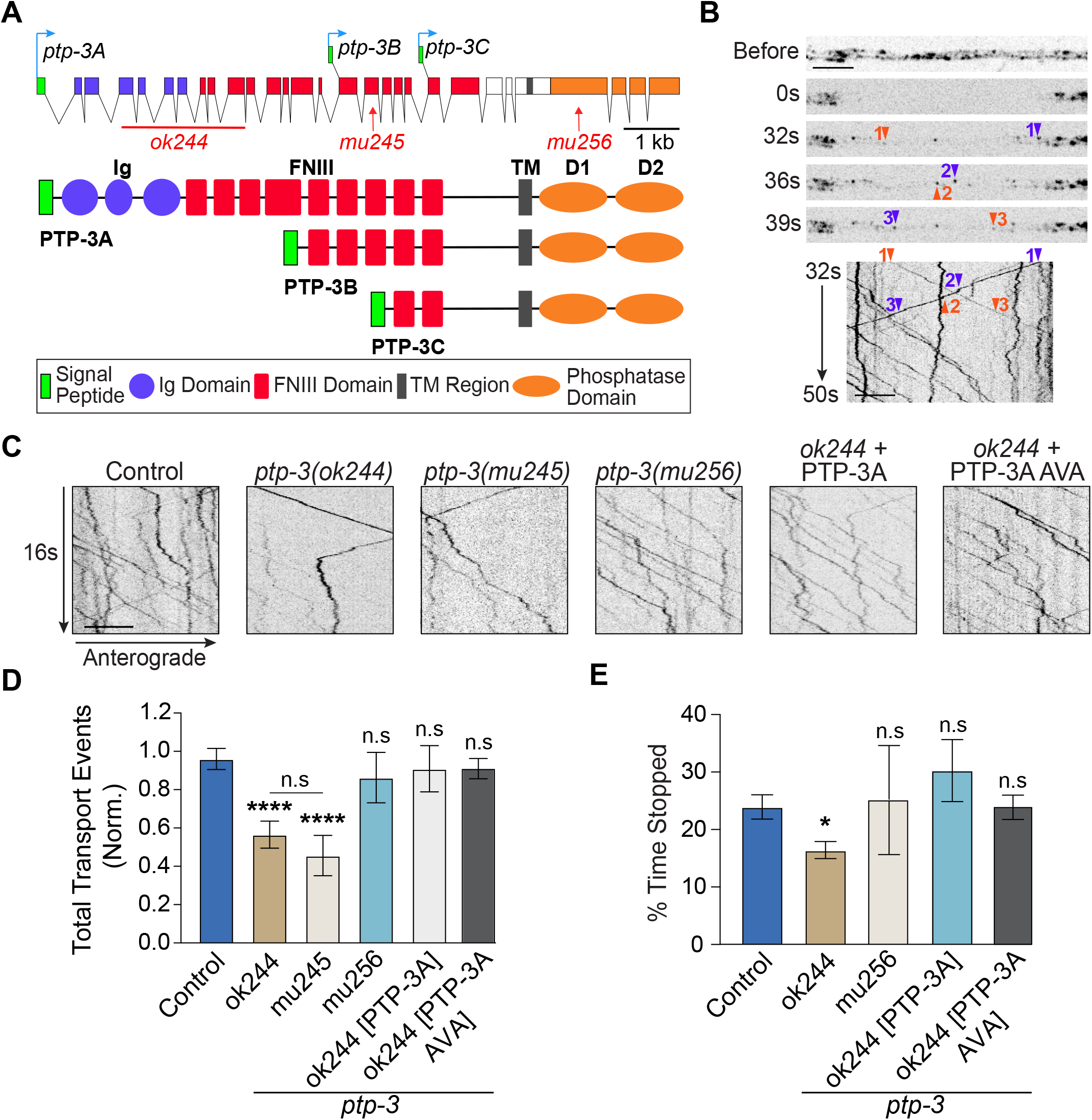
PTP-3A isoform modulates GLR-1 transport. A) Diagram of the exon/intron structure of the *ptp-3* gene (upper diagram), structure of different PTP-3 isoforms (three lower diagrams), approximate localization and nature of the alleles used in this study (also see Methods Table 1). B) Top. Sequential images and kymograph of the mCherry signal from SEP::mCherry::GLR-1 in the proximal portion of the AVA processes. The images show mCherry::GLR-1 signal prior to 1 s photobleaching and at various timepoints after showing single mCherry::GLR-1 transport vesicles (anterograde orange, retrograde purple arrowheads). Bottom. The kymograph shows all transport events from 32 – 50 s post-photobleach with time on the y-axis and position on the x-axis. The numbering of the arrowheads corresponds to the precise position and time of transport events shown in the static images above. C) Representative kymographs from the conversion of 16 s of image streaming mCherry::GLR-1 signal in the *glr-1(ky176); akIs201* background. D) Subsequent quantification of total GLR-1 transport in control (n=39) and various *ptp-3* genetic backgrounds including *ok244* (n=26)*, mu245 (n=13), mu256*(n=7) and *ok244* rescued either with PTP-3A expressed under its native promoter (n=7) or under the *flp-18* promoter (n=16). E) Kymograph quantification of the percentage of time each transport events spends stopped compared to the total time spent traveling. The number of kymographs analyzed with KymoAnalyzer were (see Methods): controls (n=15), *ok244* (n=17)*, mu256*(n=7) and *ok244* rescued either with PTP-3A expressed under its native promoter (n=5) or under the *flp-18p* promoter (n=8). Significance compared to control or as indicated: n.s.= not significant, * p < 0.05, **** p < 0.0001. All statistics used an ordinary one-way ANOVA with Dunnett’s multiple testing correction. Errors bars represent SEM and all scale bars represent 5 µm.

**Table 1:**
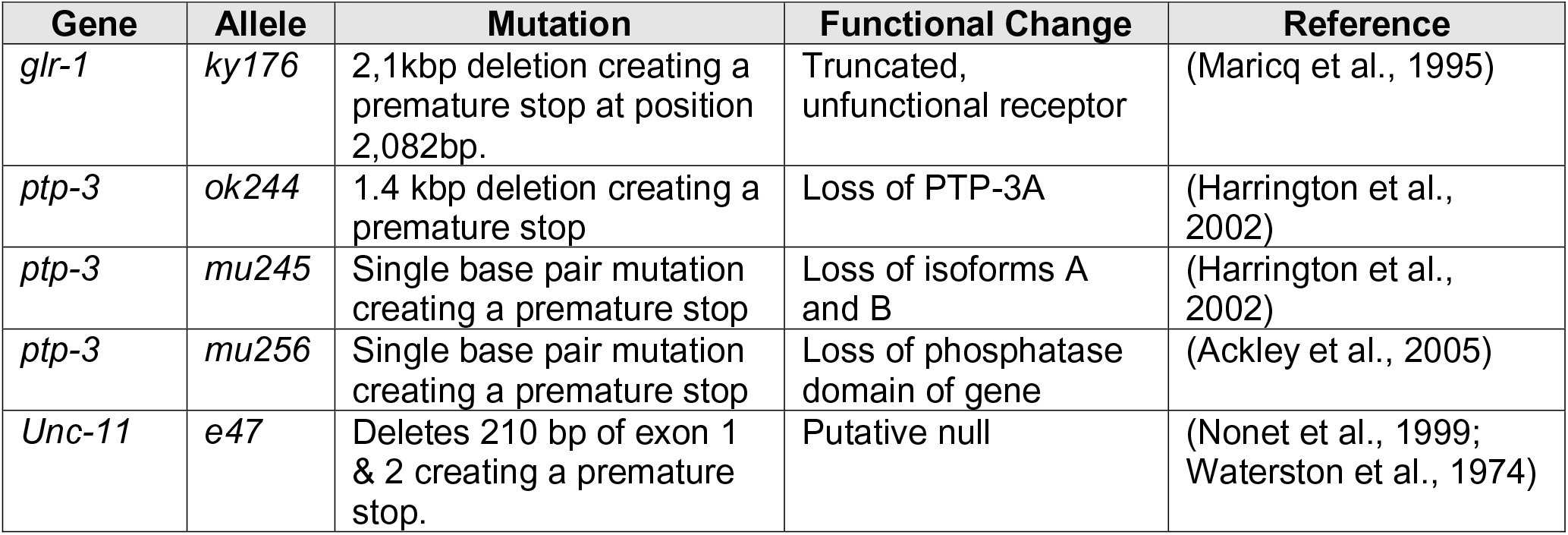
Description of the genetic alleles and reference sources.

Here, we report a postsynaptic role for the PTP-3A isoform in coordinating the transport, synaptic delivery, and synaptic recycling of GLR-1, the *C. elegans* homologue of the AMPAR subunit GluA1, which is necessary for short and long-term associative memory in *C. elegans*. By combining *in vivo,* real-time analysis of GLR-1 transport, fluorescence recovery after photobleaching (FRAP) and photoconversion with N and C-terminal cell-specific rescues, we uncovered domain-specific roles of PTP-3A in regulating synaptic localization of GLR-1. More specifically, we show that the extracellular N-terminal domain promotes GLR-1 transport from the cell body, while the intracellular C-terminal phosphatase regulates synaptic GLR-1 recycling. We link these molecular observations with specific effects on associative olfactory memory, demonstrating that changes in delivery and synaptic recycling of GLR-1 affect memory retention. We propose a model in which PTP-3A plays a dual role in regulating transport and synaptic retention of GLR-1 that is essential for long-term associative memory in *C. elegans*. Uncovering this molecular mechanism of synaptic plasticity has fundamental implications for understanding how individual neurons regulate distribution and maintenance of synaptic excitatory receptors depending on their activity in a circuit.

## RESULTS

### PTP-3A, the long isoform of PTP-3, modulates GLR-1 transport

Initial studies in rat hippocampal cell culture documented a postsynaptic specific role for LAR, PTP-δ, and PTP-σ, in recruiting GluA2 to synapses (Dunah et al., 2005; Hoogenraad et al., 2007). A local synaptic mechanism involving regulation of ß-catenin localization by LAR-RPTP was proposed (Dunah et al., 2005), but GluA2 transport was not examined. To determine how LAR-RPTPs regulate AMPAR synaptic recruitment, we started by quantifying GLR-1 transport *in vivo* in a single pair of neurons. To visualize this transport, we expressed GLR-1 tagged with the pH-sensitive Superecliptic pHluorin (SEP) and mCherry in the AVA command interneurons in the *glr-1*null background (*ky176*). We combined photobleaching and continuous imaging of mCherry::GLR-1 to observe single-vesicle GLR-1 transport events (Doser et al., 2020). The image streams were then converted into kymographs displaying lateral displacement on the x-axis and time on the y-axis which illustrate the movement of single transport events as diagonal lines (Figure 1B, orange and blue arrowheads) and immobile vesicles as vertical lines. Next, we quantified GLR-1 transport using mutations affecting the expression of specific isoforms of PTP-3, the sole *C. elegans* homologue of vertebrate LAR-RPTP. The *ok244*deletion leads to a premature stop in translation causing the loss of only the PTP-3A isoform (Figure 1A, and Methods Table 1; Ackley et al., 2005). The *mu245* mutant results in a premature stop codon that leads to loss of both PTP-3A and B isoforms (Figure 1A and Table 1; Hartin et al., 2015). The mutation in the *mu256* allele causes a frameshift and premature stop in the first intracellular phosphatase either leading to nonsense mediated decay of all mRNAs, therefore a loss of all three PTP-3 isoforms, or a truncated form of PTP-3 without phosphatase activity (Figure 1A and Table 1; Hartin et al., 2014; Ackley et al., 2005; Harrington et al., 2002). Since the isoforms are defined by differences in protein domains, such as the lack of three Ig-like domain and a few fibronectin domains in PTP-3B, the mutations affecting specific isoforms help us define how the domains affect the function of PTP-3. For example, *ok244* leads to the loss of PTP-3A, leaving only PTP-3B and 3C which lack the three Ig domains. Thus, analyzing the phenotype caused by *ok244*can reveal what the Ig domains are required for. A first look at the number of transport events (Figure 1C and 1D) shows that *ok244* and *mu245* mutants have a ∼50% decrease in GLR-1 transport events, suggesting that PTP-3A is the main isoform modulating GLR-1 transport which seems to require the Ig domains. Surprisingly, the *mu256* allele was not significantly different from controls (Figure 1C and 1D). To distinguish between a complete loss of protein translation due to the *mu256* mutation or a possible truncated version of the PTP-3 protein, we employed a reverse transcription PCR approach. Amplification of *ptp-3* from cDNA obtained from reverse transcribed mRNA with polyT or OdT primers directed at the polyA 3’UTR tail showed equal amounts of amplification for primers spanning exons 2-4 and exons 16-22, upstream of the *mu256* mutation locus, in both control and *mu256* animals (data not shown) suggesting no mRNA degradation due to *mu256*. Our data, together with previously published reports (Ackley et al., 2005; Harrington et al., 2002), suggest the phenotypes observed in *mu256* are more likely due to the presence of truncated PTP-3 rather than complete loss of all isoforms. Thus, the lack of change in GLR-1 transport observed in *mu256* suggests that PTP-3 phosphatase domains are not required for modulating GLR-1 transport. In summary, our data shows that the loss of the N-terminal Ig-like domains but not the loss of the phosphatase domains of PTP-3 is associated with a ∼50% loss of anterograde GLR-1 transport.

The decreased transport observed in the PTP-3A null, *ok244*, is rescued by expressing full length PTP-3A under the native promoter or the AVA expressing *flp-18p* promoter (Feinberg et al., 2008; Hoerndli et al., 2013) indicating a postsynaptic requirement of PTP-3A for GLR-1 transport regulation (Figure 1C and 1D). Although the amount of transport is an important determinant of GLR-1 delivery, we also recently showed that transport dynamics (i.e. transport velocity and stop frequency) play a role in GLR-1 delivery and synaptic content (Doser et al., 2020). Thus, we analyzed the percent of the time GLR-1 transport spent paused (Figure 1E) compared to the time spent moving which revealed that the *ok244* allele causes a significant decrease in stopping (Control = 24% ± 2.1; *ok244* = 16% ± 1.5). This stopping defect could be rescued by AVA-specific expression of PTP-3A (PTP-3A AVA = 25% ± 2.9). Contrary to *ok244,* the *mu256* allele did not significantly decrease stopping (*mu256* = 30% ± 5.6). We also quantified anterograde and retrograde velocities in all alleles but saw no major differences in average instantaneous velocities (Supplemental Figure 1A-D). In addition, we did not see any defects in AVA neurite length (data not shown) nor microtubule dynamics in *ok244 mutants* using the microtubule end-binding protein EBP-2 tagged with GFP (Supplemental Figure 1E-F). Together, this suggests that the decrease in GLR-1 transport observed is not due to altered processivity of molecular motors, microtubule defects, or neurite outgrowth. Furthermore, these experiments suggest that PTP-3A is important for regulating the transport of 50% of GLR-1 to synapses, which it does by stimulating both transport numbers out of the cell body and facilitating stops in dendrites which are required for synaptic delivery of receptors (Doser et al., 2020).

### PTP-3A modulates synaptic number of GLR-1 receptors

To determine whether these transport defects result in reduced synaptic content of GLR-1, we quantified fluorescence of tagged GLR-1 in the AVA process in the same *ptp-3* alleles. We used a double N-terminal tag of GLR-1 consisting of SEP (Super Ecliptic Phluorin), a pH-sensitive fluorophore that allows for quantification of GLR-1 at the synaptic surface, and mCherry to allow for the quantification of the total pool of GLR-1 receptors at synapses (Figure 2A) (Doser et al., 2020; Kennedy et al., 2010). Loss of PTP-3A (*ok244)* led to a decrease in synaptic surface GLR-1 (0.73 ± 0.06 normalized to control SEP, Figure 2C) as well as a decrease in total synaptic GLR-1 (mCherry *=* 0.66 ± 0.08 normalized to control mCherry, Figure 2D) compared to same day controls. The same analysis with *mu245* mutants showed that, like the transport phenotype, the loss of PTP-3B and 3A is the same as loss of PTP-3A alone (Supplemental Figure 2A and 2B, *mu245* normalized SEP *=* 0.75 ± 0.08, *mu245* normalized mCherry*=* 0.75 ± 0.07). The *mu256* allele showed no effect on total synaptic content of GLR-1 (Figure 2D, normalized mCherry = 1.0 ± 0.15) but decreased the SEP signal indicating a reduction in GLR-1 surface receptors (Figure 2C, normalized SEP = 0.57 ± 0.07). Decreased synaptic GLR-1 due loss of PTP-3A was rescued by expression of PTP-3A under the native promoter as well as the AVA cell-specific promoter (Figure 2B-D). This demonstrates that PTP-3A is required postsynaptically for its effect on synaptic GLR-1 trafficking. A possible explanation for decreased transport and reduced GLR-1 at synapses could be that PTP-3A is required for normal transcription and translation of GLR-1 at the soma. Alternatively, PTP-3A may be required for delivery of GLR-1 to synapses. To distinguish between these possibilities, we first measured mCherry::GLR-1 fluorescence at the AVA soma in control and *ok244.* We found no difference in fluorescence levels between genotypes (Supplemental Figure 2E-F). Second, we measured synaptic GLR-1 content in the distal regions of AVA (Figure 2E-F). Loss of PTP-3A in *ok244* showed an increase in receptors at distal synapses (SEP = 1.59 ± 0.22, mCherry *=* 1.64 ± 0.21 normalized to controls, Figure 2E-F). Together, these analyses show that there is no overall decrease in GLR-1 receptor expression but a change in distribution, which may be explained by our observations of decreased stopping of GLR-1 transport at proximal synapses when PTP-3A is lost (Figure 1E).

**Figure 2:**
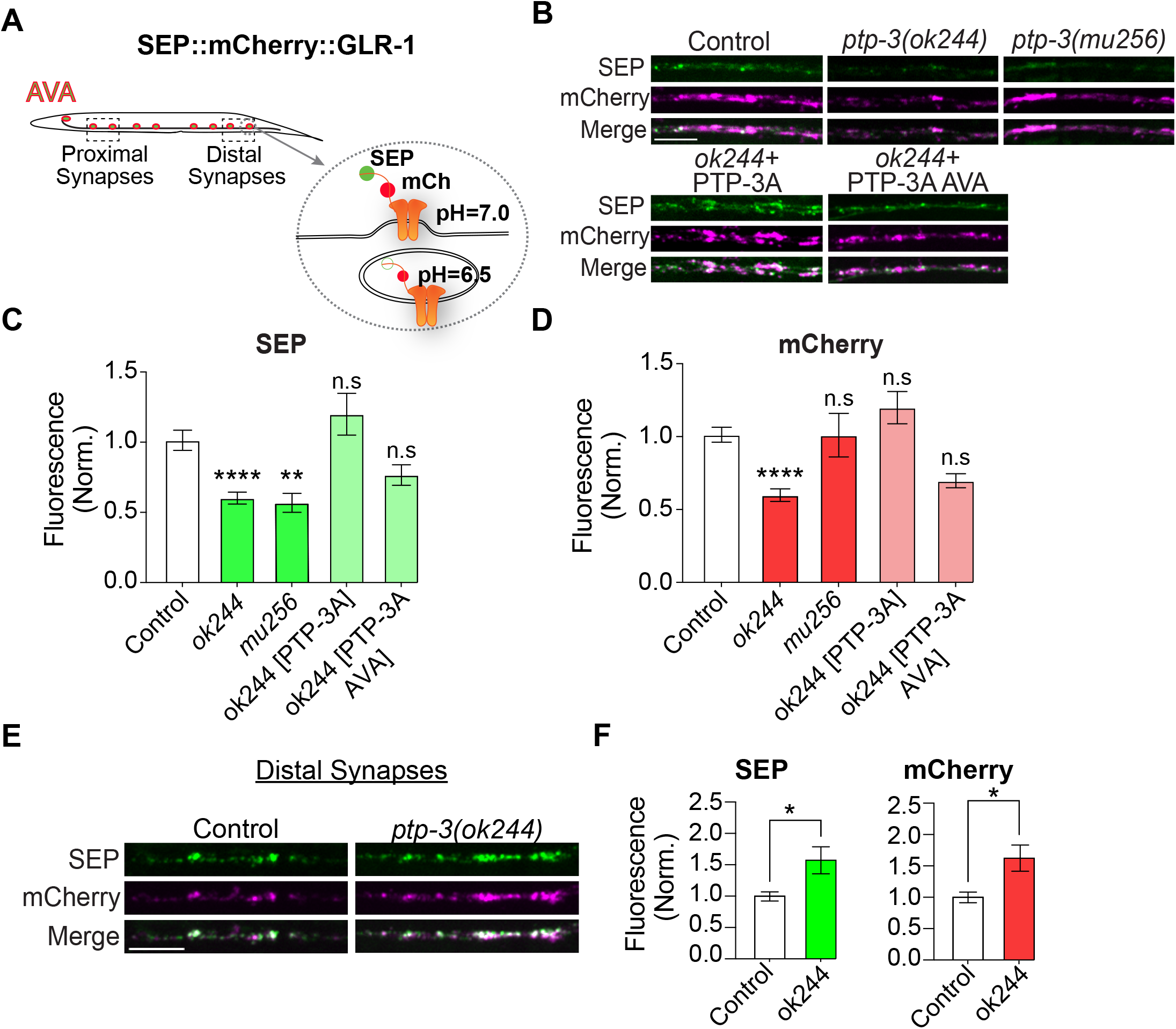
PTP-3A modulates synaptic GLR-1 levels. All worms express SEP::mCherry::GLR-1 under the *rig-3p*promoter in the *glr-1(ky176)* knockout background. Control worms represent *glr-1(ky176)* expressing SEP::mCherry::GLR-1 with no further mutations. A) Diagram of the localization of proximal and distal synapses of the AVA right and left command interneurons, with the inset showing SEP::mCherry::GLR-1 receptors fluorescing both SEP and mCherry at the synaptic surface, or only mCherry in the acidic endosome (pH 6.5). B) Confocal images of SEP and mCherry fluorescence of GLR-1 at proximal synapses. C and D) Quantification of (C) SEP fluorescence and (D) mCherry fluorescence at puncta normalized to same day controls. Control (n=56), *ok244* (n=26), *mu256* (n=18), *ok244*[PTP-3A] (n=8), *ok244* [PTP-3A AVA] (n=8). E) Confocal images of SEP and mCherry GLR-1 fluorescence in the distal synapses. F) Quantification of SEP fluorescence and mCherry GLR-1 fluorescence, normalized to same day controls. Control (n=56), *ok244* (n=22), *mu256* (n=19). Significance compared to control: n.s = not significant, * p < 0.05, ** p < 0.01, ****p < 0.0001. All statistics used an ordinary one-way ANOVA with Dunnett’s multiple testing correction. Errors bars represent SEM and all scale bars represent 5 µm.

### PTP-3A function is required in adult neurons for proper transport and synaptic GLR-1 levels

Previous work in *C. elegans* has shown a role for PTP-3 in neuronal and synaptic development (Ackley et al., 2005; Harrington et al., 2002). To address the possibility that partial loss of GLR-1 transport may be due to developmental roles of PTP-3, we analyzed transport following conditional expression of PTP-3A using an inducible heat shock approach (Hoerndli et al., 2015). Quantification of GLR-1 transport showed that heat shock treatment had no significant effect in control or *ok244* animals lacking the array (Figure 3A-C), but increased GLR-1 transport in *ok244* [*hsp16-2p::PTP-3A*] animals (Figure 3B and 3C) to nearly control levels (Figure 3A). Overall, our results suggest that PTP-3A is needed in adult neurons to modulate GLR-1 motor-dependent transport to synapses.

**Figure 3:**
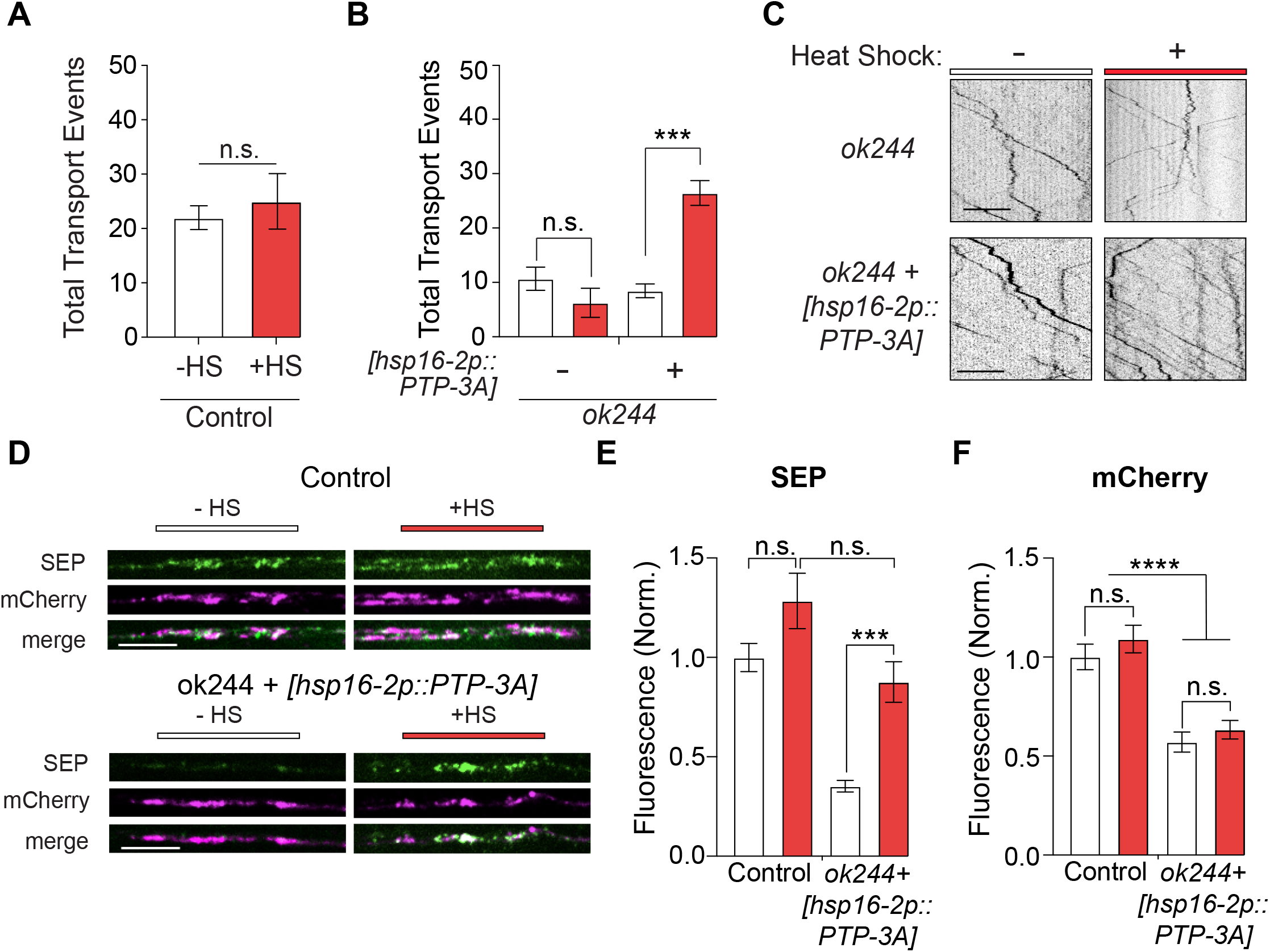
Constitutive expression of PTP-3A is required for proper transport and synaptic delivery of GLR-1 in adult animals. A-B) Quantification of the total number of GLR-1 transport events with and without 1 hr, 32° C heat shock treatment in control adult animals (A) Total transport in controls without heat shock n = 8 (white) or with heat shock n=8 (red). (B) Transport in the *ok244* background alone (-, white; n=14) or with heat-shock (-, red; n=12) as well as with *hsp16-2p::PTP-3A* with (+, white; n= 11) or without heat shock (+, red; n=16). C) Representative kymographs of GLR-1 transport events in the *ok244* background with and without the *hsp16-2p::PTP-3A* array and/or 1 hr heat shock treatment. D) Representative images of SEP and mCherry GLR-1 fluorescence in control and ok244 +_ *hsp16-2p::PTP-3A* with or without heat shock. E-F) Quantification of SEP (E) and mCherry (F) fluorescence in a proximal portion of the AVA neurite with and without 1 hr, 32° C heat shock treatment of adult animals in the *ok244* background alone (n=30 for all conditions). n.s. = not significant, * p < 0.05, ** p < 0.01, *** p < 0.001, **** p < 0.0001. All statistics used an ordinary one-way ANOVA with Dunnett’s multiple testing correction. Errors bars represent SEM and all scale bars represent 5 µm.

As mentioned previously, PTP-3 function has been shown to be important for synaptic development of the neuromuscular junction in *C. elegans* (Ackley et al., 2005; Caylor et al., 2013) but its role in synaptic development of glutamatergic command interneurons is unknown. To test whether the altered distribution of synaptic GLR-1 could be due to a role for PTP-3A in glutamatergic synapse development, we quantified the density of GLR-1 synapses (Supplemental Figures 2E and 2F), but we did not observe significant differences between *ok244* and controls. Furthermore, to determine if PTP-3A may regulate GLR-1 trafficking in adult animals regardless of its developmental role, we again used an inducible heat shock approach. However, we reasoned that rescue of synaptic GLR-1 levels would require more time than for GLR-1 transport, as observed previously (Hoerndli et al., 2015). Thus, we used two heat shock treatments separated by 12 hours to determine whether adult expression of PTP-3A was sufficient to rescue synaptic levels of GLR-1 in animals lacking PTP-3A. Our data show that heat shock had no significant effects on total dendritic or synaptic GLR-1 levels in control animals without expression of *hsp16-2::PTP-3A*. However, heat shock-induced expression of PTP-3A rescued synaptic surface GLR-1 content seen in *ok244* (Figure 3D and 3E), but almost no effect on the total pool of GLR-1 in the dendrite (as quantified by mCherry signal, Figure 3D and 3F). Together, these results show that PTP-3A has a constitutive signaling role at mature synapses.

### PTP-3A is necessary for both delivery and retention of GLR-1 at synapses

So far, our experiments show that PTP-3A can regulate GLR-1 transport and levels at synapses but it is unclear how PTP-3A does that. One hypothesis is that PTP-3A is necessary for exocytosis of GLR-1 at synapses. To test that hypothesis, we used FRAP of SEP::mCherry::GLR-1 in the same proximal region of AVA in which GLR-1 synaptic fluorescence was quantified (Figure 2 and Figure 4A). We quantified SEP and mCherry fluorescence before, immediately after, and at several timepoints following photobleaching of a section of the AVA neurite (Doser et al., 2020). Our results indicate that, compared to controls, loss of PTP-3A results in a distinctly decreased recovery rate of mCherry and SEP fluorescence (Figures 4B-E). On the other hand, the *mu256* mutants showed no effect on mCherry recovery but a strong decrease in the SEP recovery rate (Figures 4B-E). The mCherry recovery rates in these mutants are consistent with decreased GLR-1 transport observed for *ok244*and normal transport in *mu256* (Figure 1D). The decreased SEP but normal mCherry recovery rate in *mu256* indicates that this allele affects GLR-1 exocytosis or recycling rather than delivery by transport.

**Figure 4:**
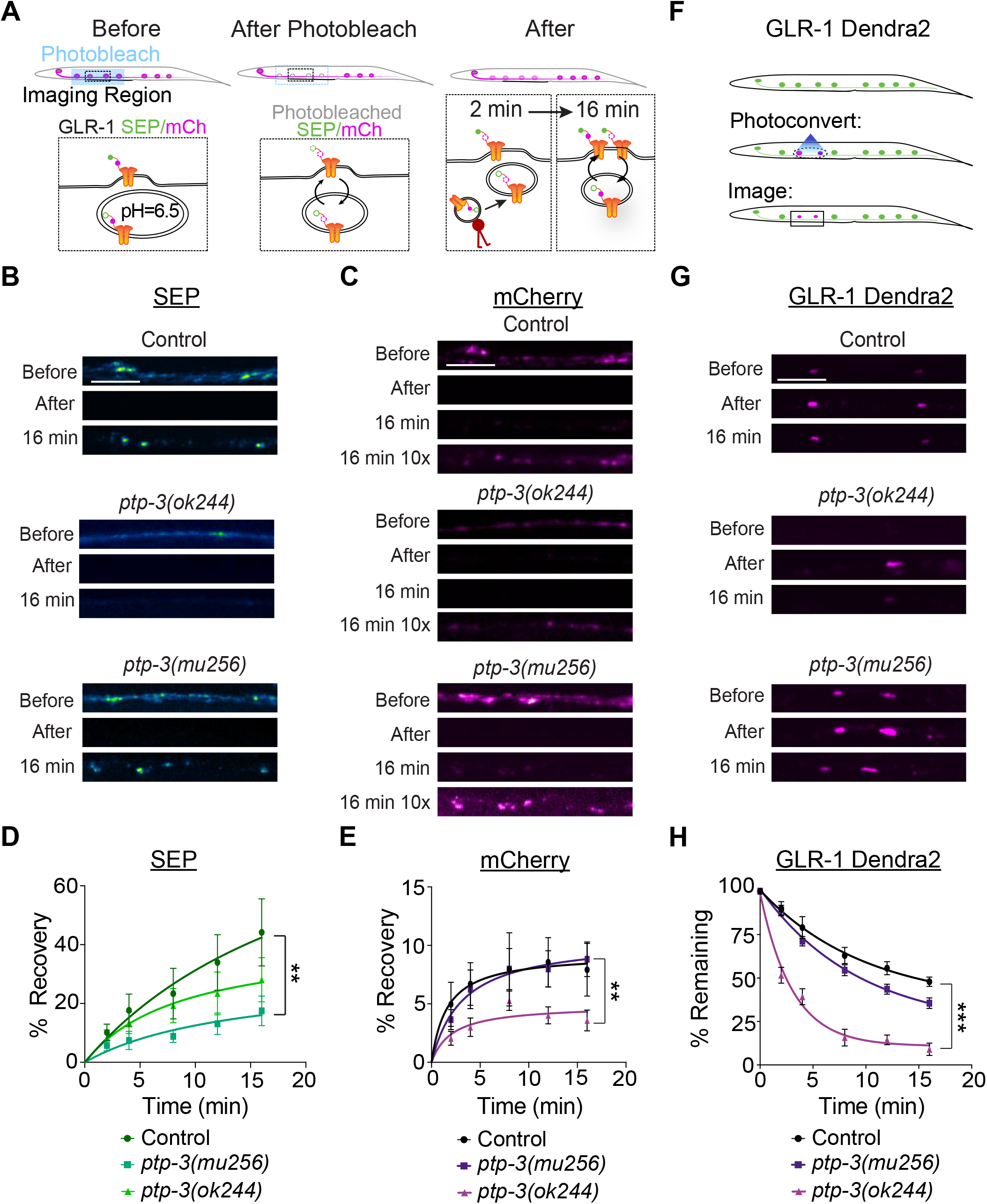
Differential regulation of delivery and removal of synaptic GLR-1 by PTP-3A. A) Cartoon illustration of the FRAP imaging location within AVA. B-C) Confocal images of (B) SEP::GLR-1 and (C) mCherry::GLR-1 taken before photobleach, directly after photobleach, 16 min post photobleach. Post processing of a 10x exposure increase of some pictures is required to see fluorescence. D-E) Quantification of the (D) SEP or (E) mCherry fluorescence recovery at 0, 2, 4, 8, 12, and 16 min post photobleaching. Control (n=7), *ok244* (n=9), *mu256* (n=9). Percent fluorescence recovery is calculated for each time point as follows: (average fluorescence (F_avg_) at time point – F_avg_ after photobleach) / (F_avg_ before). F) Cartoon illustration of the Dendra2 photoconversion protocol. G) Confocal images of GLR-1::Dendra2 in the proximal region of AVA before UV photoconversion, directly after UV photoconversion, and 16 minutes post photoconversion. H) Quantification of the red fluorescence at time points 0, 2, 4, 8, 12, and 16 min. Percent of fluorescence remaining at a time point is calculated as follows: (F_avg_ at time point - F_avg_ before) / (F_avg_ immediately after photoconversion - F_avg_ before). Control (n= 10), *ok244* (n= 10), *mu256*(n= 16) animals. ** p < 0.01, *** p < 0.001, compared to controls, using a sum of squares *F* test of the curve fit to control. Errors bars represent SEM and all scale bars represent 5 µm.

Receptor delivery to and removal from the synaptic membrane or pool affects the number of GLR-1 at synapses. To determine whether retention of GLR-1 might be affected by loss of PTP-3A or PTP-3 phosphatase domains, we used animals expressing GLR-1::Dendra2. The Dendra2 tag can be photoconverted from green to red fluorescence within a small region of the neurite (Figure 4F). Subsequent time lapse imaging of the red fluorescence provides an estimate of the removal of receptors from the synaptic pool (Hoerndli et al., 2015, 2013). *Ok244* mutant animals with a loss of PTP-3A displayed a rapid loss of GLR-1 receptors following photoconversion (only 15.7% ± 3.7 fluorescence remaining after 16 minutes compared to 63 % ± 4.6 in control animals, Figure 4G and 4H). In *mu256* mutants that lack PTP-3 phosphatase domains, loss of GLR-1 receptors was also faster than controls, but much slower than *ok244* (54.5 % ± 3.5 after 8 minutes, Figure 4G and 4H). Indeed, when we quantified the number of remaining photoconverted receptors still present after 2 hours, both *ok244* and *mu256* had significantly fewer receptors remaining compared to controls, but not significantly different from each other (Supplemental Figure 3A).

Since there is both enhanced removal and decreased delivery rates in mutants without PTP-3A, it is not clear which of these two mechanisms contributes more to decreased synaptic GLR-1 numbers. To try to address this, we used a genetic loss of function of UNC-11, the *C. elegan*s homologue of a clathrin adaptor protein AP180, necessary for the endocytosis of GLR-1 (Burbea et al., 2002). We made double mutants with *unc-11* loss-of-function *(lf)* and *ptp-3(ok244)* containing SEP::mCherry::GLR-1 (Supplemental Figure 3C-and 3D). Quantification of synaptic SEP and mCherry signals in *unc-11(lf)* showed that a lack of endocytosis led to higher levels of SEP::GLR-1 compared to controls (Supplemental Figure 3B-C) but did not affect mCherry::GLR-1 signal. This indicates that loss of *unc-11* clearly affected synaptic surface levels of GLR-1. Here we can make a few predictions. First, if loss of PTP-3A affects synaptic GLR-1 by acting on delivery and exocytosis, then double mutants with *unc-11(lf)* would have less synaptic SEP::GLR-1 signal than *unc-11(lf)* alone. Second, if delivery and exocytosis defects are secondary to increased removal, then double mutants with *unc-11(lf)* would appear to have normal or increased SEP::GLR-1. Our results indicate that *ptp-3A(ok244); unc-11(lf)* show increased SEP::GLR-1 compared to controls and *unc-11(lf)* (Supplemental Figure 3C). In addition, total levels of mCherry::GLR-1 are indistinguishable between controls, *unc-11(lf)* and *unc-11(lf); ptp-3A(ok244)* doubles (Supplemental Figure 3D). Altogether, these experiments suggest that removal is the main mechanism by which PTP-3A regulates retention of GLR-1 receptors at the synaptic surface.

### Differential roles of PTP-3A N-terminal and C-terminal domains in regulating GLR-1 transport and synaptic retention

Since our analyses suggest that loss of PTP-3A is equivalent to the loss of both the A and B isoform, which differ by the presence of the N-terminal Ig-like domains and a few fibronectins, we hypothesized that the N-terminal of PTP-3A is specifically important for normal GLR-1 transport, delivery, and retention. To test this hypothesis, we engineered a construct consisting of the membrane insertion signal peptide and the two most N-terminal Ig-Like domains of PTP-3A by AVA neurons (Figure 5B) and expressed it in *ptp-3(ok244)*. We first quantified GLR-1 transport in controls, *ok244,* and *ok244* with the N-terminal constructs (Figure 5A-C). Interestingly, postsynaptic expression of the PTP-3A N-terminal domain rescued GLR-1 transport compared to *ok244* alone (Figure 5C, all values normalized to wild-type controls, *ok244 +* N-term 0.863 ± 0.066, *ok244* alone 0.595 ± 0.074). In addition, we also expressed the C-terminal domain with the transmembrane domain (Figure 5B) in *mu256* mutants only in AVA neurons but did not observe any change in GLR-1 transport (Figure 5D). To determine if the C-terminal domain of PTP-3A has a function independent of the rest of the protein, we expressed it in *ok244* mutants but saw no rescue of GLR-1 transport (data not shown). Altogether these transport analyses are consistent with the N-terminal but not C-terminal of PTP-3A being a modulator of GLR-1 transport numbers.

**Figure 5:**
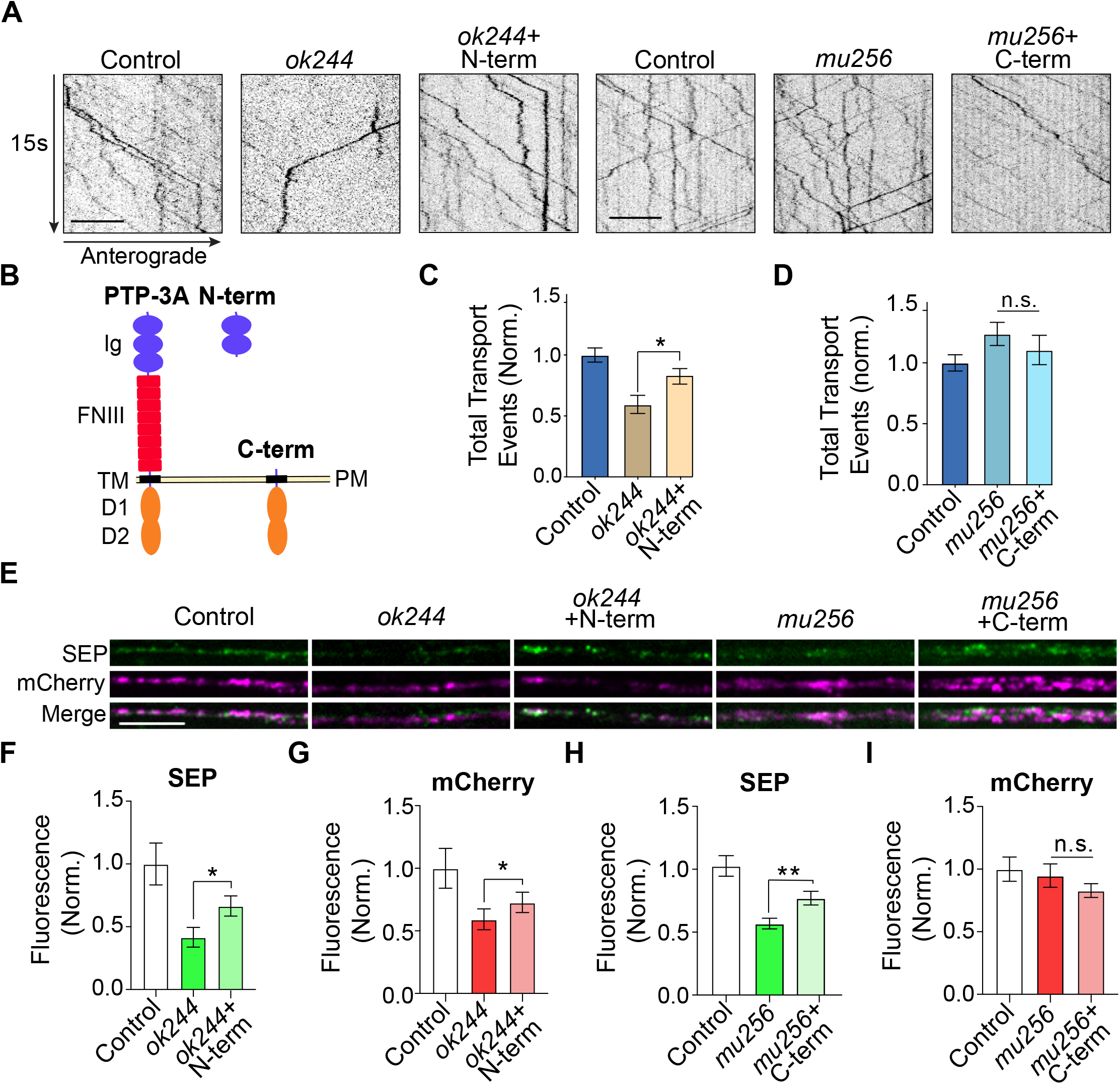
The N-terminal and C-terminal domain of PTP-3A regulate different aspects of GLR-1 transport to synapses. A) 15 s of representative kymographs from each group. ‘N-term’ represents the two most terminal Ig-like domains of PTP-3A expressed under the AVA specific *flp-18p* promoter in the *ok244* background. ‘C-term’ represents the transmembrane domains and internal C-terminus of PTP-3A including the dual-phosphatase domain expressed under the AVA specific *flp-18p* promoter in the *mu256* background. All constructs retain the signal peptide (not shown). B) Cartoon schematic of the N- and C term constructs. C) Quantification of GLR-1 transport in *ok244* and *ok244* + N-term (normalized to control). Control (n=25), *ok244* (n=26), *ok244* + N-term (n=16); control (n=26), *mu256* (n=10), *mu256* + C-term (n=10). D) Quantification of GLR-1 transport in *mu256* and *mu256*+ C-term (normalized to controls). E) Confocal images of SEP::GLR-1 and mCherry::GLR-1 from each group. F-G) Quantification of (F) SEP and (G) mCherry::GLR-1 fluorescence normalized to control. Control (n=15), *ok244* (n=11), *ok244* + N term (n=24). H-I) Quantification of (H) SEP fluorescence and (I) mCherry GLR-1 fluorescence normalized to controls (n=15), *mu256* (n=16), *mu256* + C term (n=18). n.s. = not significant, * p < 0.05, ** p < 0.01. Statistical comparisons used an unpaired Students’ T-test. Errors bars represent SEM and all scale bars represent 5µm.

Our previous analyses of GLR-1 synaptic content (Figure 2) showed that loss of PTP-3A led to changes in both total and synaptic numbers of surface GLR-1, whereas loss of the phosphatase domains only affect synaptic numbers of surface GLR-1. Here we tested whether expression of PTP-3A N-terminal domain in AVA rescued both total and synaptic surface levels of GLR-1. Our data indicate that N-terminal PTP-3A expression in *ok244* in AVA significantly rescues transport, total synaptic and surface numbers of GLR-1 (Figure 5E-G). We then tested whether expression of the C-terminal domain of PTP-3A was sufficient to rescue the lower levels of GLR-1 receptors observed for *mu256* (Figure 2). Our data show that indeed C-terminal expression is sufficient to partially rescue the number of synaptic GLR-1 surface receptors (Figure 5E, H, I). Taken together, these results further suggest a postsynaptic role for the N-terminal domain of PTP-3A in regulating transport and synaptic delivery of GLR-1, whereas the C-terminal of PTP-3A helps retain GLR-1 at the synaptic surface.

### PTP-3A is necessary for olfactory associative memory but not learning

Several previous studies have shown that decreases in synaptic GLR-1 numbers are associated with defects in associative olfactory memory in *C. elegans* (Hadziselimovic et al., 2014; Stetak et al., 2009; Vukojevic et al., 2020, 2012). To determine if synaptic GLR-1 defects due to loss of PTP-3A are associated with learning and memory, we used population-based aversive olfactory learning and memory assays (Bargmann et al., 1993; Hoerndli et al., 2009; Stetak et al., 2009; Vukojevic et al., 2012). We quantified olfactory attraction to 0.1% diacetyl, a natural attractant for *C. elegans*, in naïve animals and in animals that underwent diacetyl-coupled starvation training (Figure 6A and 6B). We observed that loss of PTP-3A (*ok244)* or all phosphatase domains (*mu256)* did not affect naïve attraction, nor did it prevent learning the negative association of diacetyl with starvation (Figure 6B). Analysis of diacetyl attraction after one hour of conditioning and one hour of recovery, revealed effects on <u>S</u>hort-<u>T</u>erm <u>A</u>ssociative <u>M</u>emory (STAM) (Stetak et al., 2009). <u>Lo</u>ng-Term <u>A</u>ssociative <u>M</u>emory (LTAM) was assayed by repeating negative association with 30 minutes of recovery in between and then quantifying attraction 24 hours after training (Figure 6A and 6C) (Vukojevic et al., 2012). These memory assays revealed that loss of PTP-3A significantly decreases STAM and LTAM whereas loss of the phosphatase domains only significantly impaired LTAM (Figure 6B and 6C). To determine if this effect on STAM and LTAM required pre- or postsynaptic expression of PTP-3A, we repeated the memory assays with *ok244* mutants expressing PTP-3A under an AVA-specific promoter (Figure 6D and 6E). Rescue of PTP-3A expression in AVA alone rescued both STAM (1h memory) and LTAM (24h memory) defects observed in animals lacking PTP-3A (Figure 6D and 6E), without altering either naïve attraction or learning. These data show that postsynaptic expression of PTP-3A is essential for both short and long-term olfactory associative memory.

**Figure 6:**
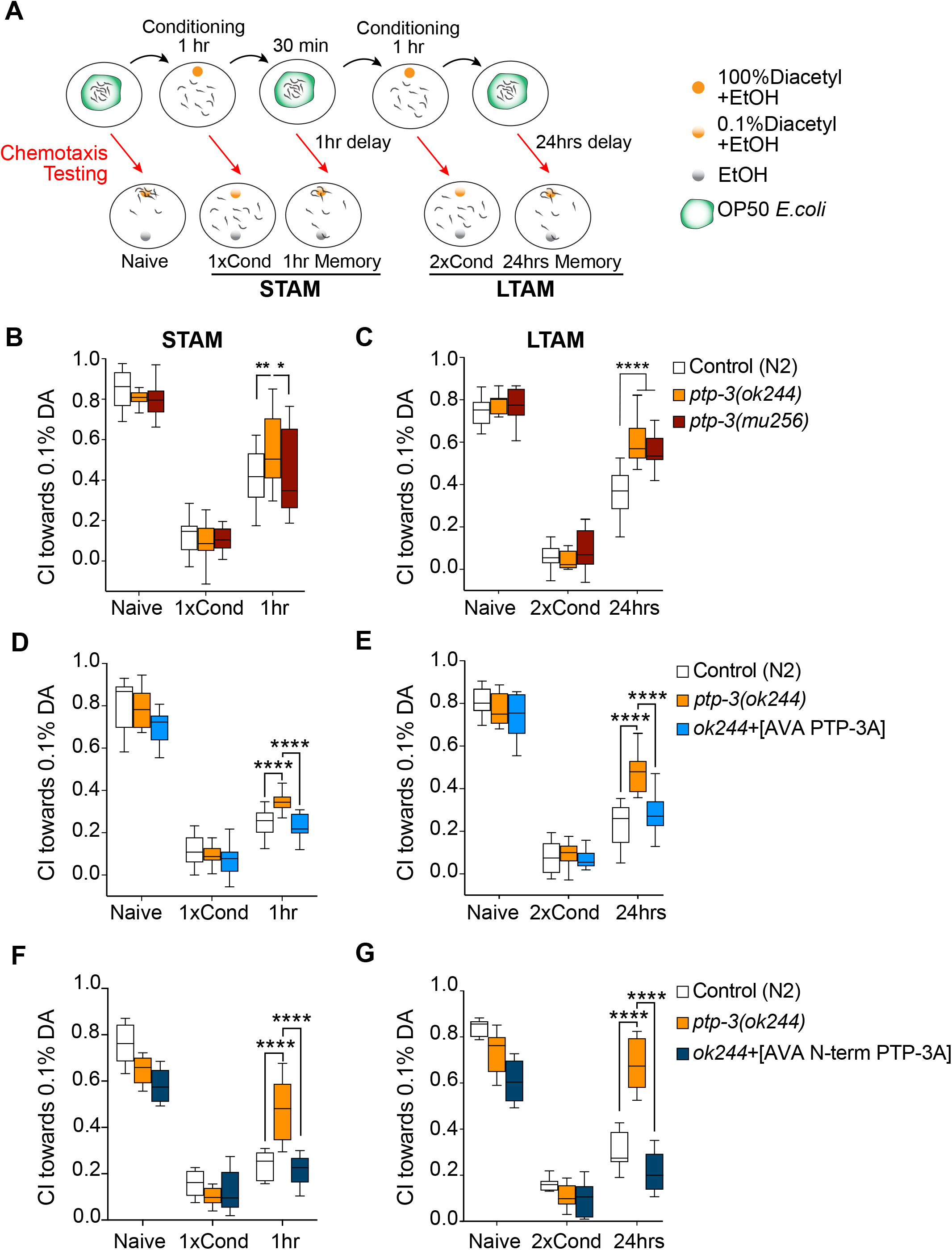
Loss of PTP-3A function affects short and long-term associative memory. A) Illustration of the chemotaxis associative memory protocols for **<u>S</u>**hort-**<u>T</u>**erm **<u>A</u>**ssociative **<u>M</u>**emory (STAM) and **<u>L</u>**ong-**<u>T</u>**erm **<u>A</u>**ssociative **<u>M</u>**emory (LTAM). B) Quantification of the chemotaxis towards 0.1% Diacetyl of unconditioned worms (naïve), worms conditioned for 1 hour with 1% Diacetyl (DA) without food (1xCond) and animals conditioned but transferred to food for 1 hour (1hr). The following genotypes were tested: control (N2) (white bars, N=18 trials), *ptp-3(ok244)* (orange bars, N=15 trials) and *ptp-3(mu256)* (red bars, N=12 trials). C) Tukey plot of the quantification of the chemotaxis towards 0.1% Diacetyl of unconditioned worms (naïve), worms conditioned twice for 1 hour with 1% Diacetyl (DA) without food with 30 minutes rest between conditioning (2xCond) and animals conditioned but transferred to food for 24 hours (24hrs). The following genotypes were tested: control (N2) (white bars, N=18 trials), *ptp- 3(ok244)* (orange bars, N=15 trials) and *ptp-3(mu256)* (red bars, N=12 trials). D-E) Tukey plot of the quantification of STAM (D) and LTAM (E), in control (N2, white), *ptp-3(ok244*) (orange), and *ptp-3(ok244)* expressing PTP-3A (blue) in AVA. N=15 trials for each group. F-G) Tukey plot of the quantification of STAM (F) and LTAM in (G) in control (N2, white), *ptp-3(ok244)* (orange) and *ptp-3(ok244)* with AVA expression of the N-term 2 Ig-like domains (burgundy) (N=12 trials for STAM and N=9 trial for LTAM for all genotypes). Chemotaxis Index (CI) = (worms at DA – worms at EtOH) / total number of worms. * p < 0.05, ** p < 0.01, *** p < 0.001, **** p < 0.0001, using a 2-way ANOVA with Bonferroni correction for multiple testing. Tukey plots show whiskers as min and max, box length as interquartile range and middle line as median.

Our previous analyses of synaptic GLR-1 numbers and transport dynamics in animals lacking PTP-3A correlated with functional consequences in memory retention (Figure 6B-E). To test whether this holds true for the N-terminal AVA expression of PTP-3A, we tested both STAM and LTAM in controls, *ok244,*and *ok244* with AVA expression of the N-terminal of PTP-3A containing the Ig-like domains. Consistent with our previous analyses, animals lacking PTP-3A (*ok244*) exhibited defects in STAM and LTAM without any defects in learning compared to controls (Figure 6F and 6G). AVA expression of the Ig-like domains completely rescued these defects back to control level. Altogether, our experiments not only uncover a new role for PTP-3A in regulating long-distance GLR-1 transport, delivery and synaptic retention but also identify domain specific functions of PTP-3A. Our results suggest a hypothetical model (Figure 7) in which the N-terminal domain of PTP-3A regulates the amount GLR-1 transport and receptor delivery to synapses whereas the C-terminal phosphatase domains regulate local synaptic retention of GLR-1.

**Figure 7:**
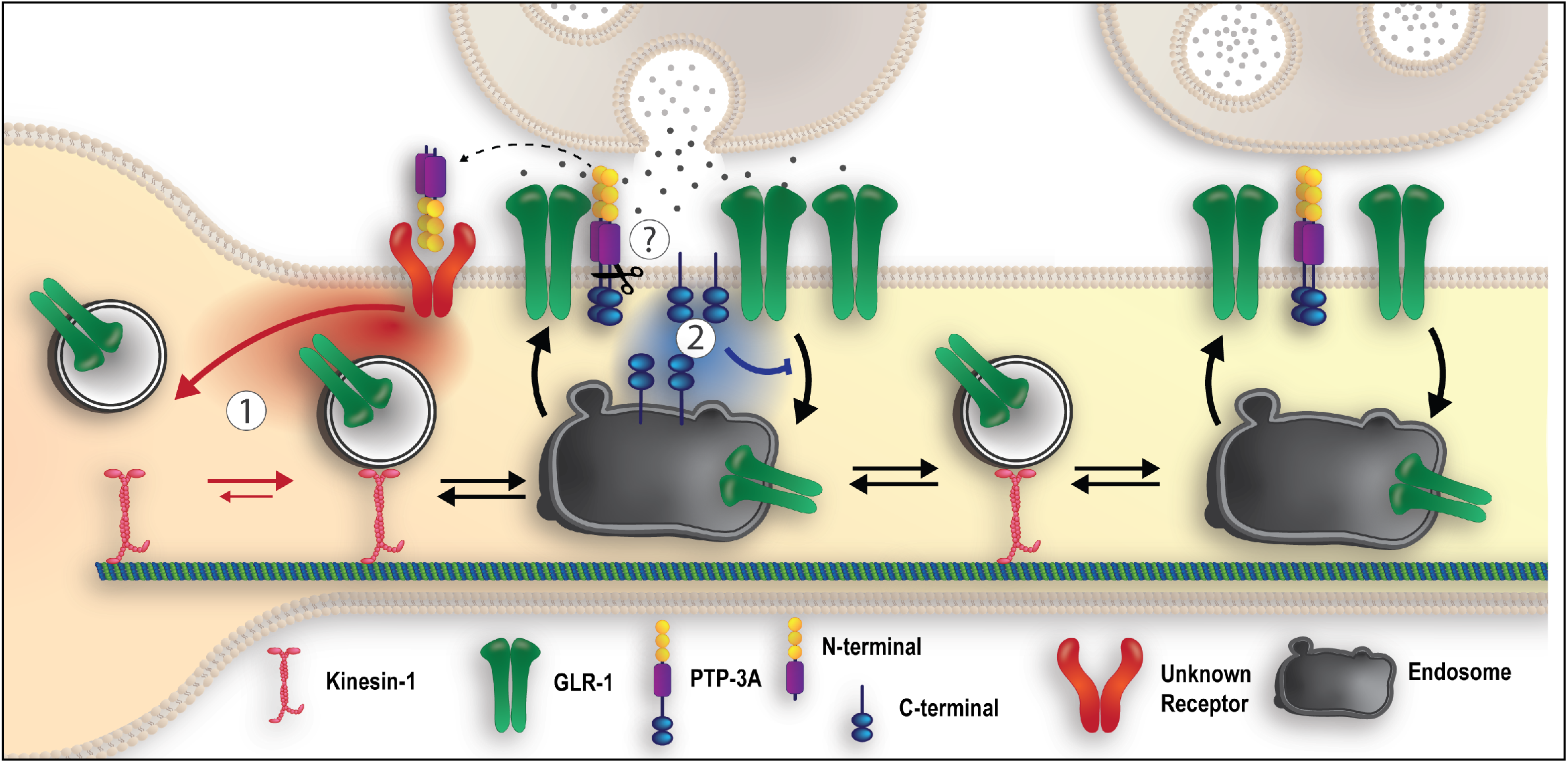
Hypothetical model of the dual function of PTP-3A in regulating GLR-1 transport and synaptic retention. Synaptic activity and glutamate release may lead to cleavage of the extracellular and intracellular domains of PTP-3A **(?)**. The N-terminal domains could bind to an unknown receptor to promotes GLR-1 transport **(1)**. The cleaved C-terminal with the phosphatase domains either remains at the synaptic surface or is internalized **(2)** where it can dephosphorylate unidentified substrates leading to retention of GLR-1 at synapses.

## DISCUSSION

Several key studies have now shown that activity regulated molecular motor-dependent transport of AMPARs is an evolutionarily conserved mechanism that is essential for synaptic function and plasticity (Gutiérrez et al., 2021; Hangen et al., 2018; Hoerndli et al., 2015). Although these studies present evidence that this transport system is necessary and that global transport levels are regulated by neuronal activity, they have not identified the regulatory mechanisms of AMPAR delivery to synapses nor how neurons integrate synaptic demands to achieve proper homeostatic balance.

Here, we present evidence that the PTP-3A isoform, the closest *C. elegans* homologue of vertebrate LAR-RPTP (Ackley et al., 2005; Harrington et al., 2002), is a key signaling component that enables the coordinated distribution of synaptic GLR-1/AMPAR. By analyzing long-distance transport and local synaptic dynamics of GLR-1, we determined that PTP-3A modulates global trafficking and synaptic retention of AMPARs. Our data show that two separate domains of the transmembrane PTP-3A differentially regulate GLR-1 transport. The N-terminal extracellular portion containing only Ig-like domains of PTP-3A regulates the number of transport vesicles containing GLR-1 that originate from the soma, which in turn impacts delivery of receptors to synapses. This process is dependent on the presence of the two most N-terminal Ig-like domains of PTP-3A. The C-terminal intracellular portion of PTP-3A containing two phosphatase domains regulates synaptic retention of GLR-1 by acting locally on receptor delivery and retention. In addition, we show that postsynaptic PTP-3A functions are necessary for proper associative olfactory memory that depend on glutamatergic neurotransmission in *C. elegans* command interneurons

### Cell-specific regulation of GLR-1/AMPAR transport by PTP-3A/LAR-RPTP

Previous studies in *C. elegans* have shown a presynaptic role for PTP-3A working with *syd-2*/Liprin-α and Nidogen-1 at the neuromuscular junction during synapse development (Ackley et al., 2005; Hartin et al., 2015). In addition, these *C. elegans* studies indicated a neurodevelopmental role for PTP-3 (Ackley et al., 2005; Harrington et al., 2002) similar to the LAR-RPTPs homologue in vertebrates (Coles et al., 2015; Won and Kim, 2018). More specifically, it was observed that the PTP-3B isoform was necessary for axon guidance whereas the PTP-3A was important for regulating synapse formation. Our data show that cell-specific expression of PTP-3A in AVA is sufficient to rescue transport and synaptic content of GLR-1, suggesting a postsynaptic function of PTP-3A in GLR-1 regulation at excitatory synapses. Furthermore, synaptic GLR-1 levels could be rescued by acute expression of PTP-3A (Figure 3) indicating that PTP-3A has more of a constitutive role in GLR-1 transport, delivery, and retention at the synapse.

A few key studies in dissociated rat hippocampal culture have shown a postsynaptic role for LAR-RPTPs in AMPAR retention, particularly of GluA2 retention (Dunah et al., 2005; Uetani, 2000; Wyszynski et al., 2002). The proposed mechanism by which LAR-RPTPs increase retention of AMPARs was based solely on a local synaptic role for LAR-RPTPs in dephosphorylating proximal proteins recruited by Liprin−α. In this study we also show a postsynaptic role for PTP-3/LAR-RPTP, but we add new observations that show that PTP-3/LAR-RPTP is important not only for synaptic retention of GLR-1/AMPAR but also for modulating long-distance transport. Two sets of experiments suggest that the extracellular domain of PTP-3A is specifically important outside of the synapse for normal GLR-1 transport. First, the *mu256* allele that lacks only the C-terminus did not alter GLR-1 transport (Figure 1). Second, cell-specific expression of the two most N-terminal Ig domains of PTP-3A rescued GLR-1-transport (Figure 5).

These results suggest that the N-terminal extracellular domain of PTP-3A has a signaling function, and perhaps interacts with a membrane-bound signaling protein or receptor that promotes AMPAR transport and synaptic delivery. Shedding of the extracellular domain of LAR-RPTPs has been documented in non-neuronal cell culture experiments (Serra-Pagès et al., 1995; Streuli et al., 1992) and shown to be regulated by cytoplasmic calcium and PKC signaling (Aicher et al., 1997). Cell adhesion molecules of the Immunoglobulin superfamily, such as NCAM1 and NCAM2, undergo cleaveage (Kim et al., 2021). In addition, cleavage and shedding of the extracellular region of neuroligins and neurexins have been reported to have signaling roles *in vivo* in vertebrate models (Borcel et al., 2016; Trotter et al., 2019; Venkatesh et al., 2017, 2015). However, a signaling role for the shed ectodomain has not been determined for LAR-RPTP nor PTP-3A. Altogether, our studies of *in vivo* transport of GLR-1 in *C. elegans* reveal the exciting possibility that LAR-RPTPs could be modulating levels of AMPAR transport by a mechanism akin to autocrine signaling involving cleavage of the extracellular domain of LAR-RPTP.

### Regulation of local synaptic GLR-1 levels by PTP-3A/LAR-RPTP

As mentioned above, a few studies have documented postsynaptic localization and roles for LAR-RPTPs in regulating synaptic retention of AMPARs (Dunah et al., 2005; Hoogenraad et al., 2007). These studies suggested that LAR-RPTPs recruit and aid the retention of AMPARs at synapses via direct interactions and dephosphorylation of local synaptic proteins. More recent *in vivo* studies using triple conditional knock-out of LAR, PTP-δ and PTP-σ models on the other have shown no effect on either excitatory or inhibitory synaptic function in the CA-1/3 neurons of the hippocampus in mice (Emperador-Melero et al., 2021; Sclip and Südhof, 2020). Our results show that PTP-3A is acting on two levels of the AMPAR trafficking process. The first pertains to promoting transport of AMPARs as discussed above. The second pertains to the regulation of AMPAR synaptic retention. Our data show that loss of PTP-3A results in substantial decreases in total synaptic and synaptic surface numbers of GLR-1 (Figure 2) without affecting the density of synapse numbers (Supplemental Figure 2C-D). In addition, cell specific rescue of PTP-3A supports a postsynaptic, and likely local, synaptic role for PTP-3A in regulating synaptic GLR-1 numbers. Using FRAP and Dendra2 photoconversion approaches to analyze delivery and removal of synaptic GLR-1 in *ptp-3* mutants (Figure 4) and clathrin adaptor mutants (*unc-11(lf),* Supplemental Figure 3A-B), we further show that the decrease in GLR-1 at the synaptic surface is due to increased removal of GLR-1, which requires endocytosis (Supplemental Figure 3C-D). Finally, by comparing FRAP of GLR-1 in strains lacking PTP-3A (*ok244*) or the PTP-3 phosphatase domains (*mu256)*, we present evidence for a role of the D1 and D2 phosphatase domains in synaptic surface retention of GLR-1. Altogether, these data are consistent with a model in which PTP-3A phosphatase domains are important at synapses to maintain a functional number of GLR-1 receptors.

The results could be mediated by direct dephosphorylation of downstream targets of PTP-3A or interactors which bind to the D2 domain of PTP-3A similar to what has been observed for vertebrate LAR-RPTPs (Dunah et al., 2005; Serra-Pagès et al., 1998; Wyszynski et al., 2002). Genetic evidence presented previously for binding and synaptic co-localization of PTP-3 and SYD-2 (the *C. elegans* homologue of Liprin-α; Ackley et al., 2005) mirrors the vertebrate binding and synaptic localization of LAR-RPTP and Liprin-α (Dunah et al., 2005; Hoogenraad et al., 2007). In vertebrates, LAR, Liprin-α, and GRIP-1 form a complex (Wyszynski et al., 2002); however, it is unclear if a similar complex is present in *C. elegans* as a homologue of GRIP-1 in *C. elegans* has not been reported. Since a loss of PTP-3 can lead to mislocalization of SYD-2, it is possible that the decrease in synaptic GLR-1 levels in PTP-3A mutants could be due to a lack of synaptic SYD-2 rather than a direct signaling role of PTP-3A at synapses. Finally, in addition to Liprin-α, a Rho/Rac guanine nucleotide exchange factor named Trio can interact with the D2 domain of LAR-RPTP (Debant et al., 1996) and has been reported to play a critical role in synaptic plasticity (Herring and Nicoll, 2016). An interaction between PTP-3 and the *C. elegans* homologue of Trio, UNC-73 (Williams et al., 2007), has not yet been assessed.

Potential substrates of PTP-3-mediated tyrosine dephosphorylation in *C. elegans* neurons have been identified, including several members of the mitogen activated kinase family (MAPK), multiple tyrosine kinases (FER, MET and NTRK2), and cyclin-dependent kinase 5 (CDK5) (Mitchell et al., 2016). Interestingly, previous studies have shown that synaptic levels of GLR-1 are decreased in a *cdk-5* loss-of-function mutants (Juo et al., 2007). In addition, this study showed that CDK-5 interacted with the clathrin adaptor, AP180, which regulates GLR-1 endocytosis. CDK-5 is a potential substrate for PTP-3 and we found that recycling of GLR-1 is also perturbed in PTP-3A loss-of-function mutants making CDK-5 an interesting potential target and effector in this signaling pathway. Although classical CDKs are usually inactivated by phosphorylation (Shah and Rossie, 2018), the role of phosphorylation in the activation of CDK-5 in neurons remains unclear in vertebrates (Kobayashi et al., 2014) and unknown in *C. elegans*.

The D1and D2 domains of PTP-3A are highly conserved compared to the D1 and D2 domains of LAR-RPTP. The D1 domain in LAR-RPTP has many downstream targets including β-catenin which has been shown to co-immunoprecipitate with GluA2, Liprin-α, and GRIP-1 (Dunah et al., 2005; Hoogenraad et al., 2005; Wyszynski et al., 2002). GluA2 tyrosine phosphorylation is associated with its own internalization (Han L. Tan et al., 2020; Yong et al., 2020), but it remains to be shown whether GluA2 and GluA1 can be dephosphorylated by LAR/PTP-3A. Taken together, PTP-3A could be affecting local GLR-1 retention through interactions with scaffolds or dephosphorylation of key downstream targets, including AMPAR subunits.

Although intensely studied in the last 25 years, the regulation of LAR-RPTP activation is not well understood and has been majorly focused on the role of the extracellular domains acting as ligands for trans-synaptic partners. Overall, it is unclear whether LAR-RPTPs mediate their effect through binding at their D2 domains or through activity of the phosphatase D1 domain. In addition, how activity of the phosphatase domains is regulated is not well understood. Early models predicted that de-clustering of LAR-RPTPs led to loss of phosphatase activity (Serra-Pages et al., 1994), but more recent structural studies suggest that tight clustering of LAR-RPTP could inhibit phosphatase activity since D1 domains can form tight homophilic interactions (Xie et al., 2020). In this latest model, it is possible that cleavage of LAR-RPTP could lead to decreased clustering and increased activity of the D1 domains. In this context, it is interesting to note that PKC activation and increased calcium influx through calcium channels lead to cleavage, shedding of the extracellular domain and then internalization of LAR-RPTP (Aicher et al., 1997). In our model (Figure 7), the D1 and D2 domains would be facing the cytoplasmic side and thus able to interact with substrates for dephosphorylation. Since it is known that synaptic activity leads to a local increase in PKC activity (Callender and Newton, 2017; Hirbec et al., 2003) and lysosomal exocytosis causing release of metalloproteases (Padamsey et al., 2016), it is tempting to speculate that synaptic activity might lead to cleavage of the extracellular domain and perhaps internalization of intracellular domains of PTP-3A as shown Figure 7. At the very least, this provides an interesting model to test for future studies.

### How AMPAR transport and synaptic retention affects learning and memory

Although it is now clear that the number and function of AMPARs at synapses can be directly associated with synaptic plasticity and behavioral learning and memory (Groc and Choquet, 2020; Tan et al., 2020; Volk et al., 2015), the role of long-distance transport in learning and memory has not been well studied. A logical assumption would be that any process leading to a decreased content of receptors at excitatory synapses would affect induction of synaptic plasticity or learning, whereas any process affecting maintenance of receptors after plasticity would affect memory. The command interneuron AVA plays a special role in olfactory associative memory in *C. elegans* (Hendricks et al., 2012; Stetak et al., 2009; Vukojevic et al., 2012). The most recent studies to date show that activity-dependent regulation of AMPAR transport is necessary for synaptic plasticity (Hangen et al., 2018; Hoerndli et al., 2015). However, we do not yet understand how this relates to learning and memory. Many studies have validated how synaptic plasticity relates to behavioral learning and memory (Huganir and Nicoll, 2013) but it is unclear how necessary transport, delivery and removal are for learning and/or memory. Here we show that both short and long-term associative memory in *C. elegans* depend on modulation of GLR-1 transport and synaptic retention by the PTP-3A isoform (Figure 6).

### A model for coordination of AMPAR transport and synaptic maintenance

Our analysis of transport, FRAP, synaptic imaging and behavior has enabled us to identify two domains of PTP-3A with differential effects on GLR-1 trafficking. We propose a model in which this dual function regulates the distribution of AMPARs to synapses. More specifically, we hypothesize that cleavage of PTP-3A promotes transport through the N-terminal autocrine effect, and synaptic retention through local liberation of the C-terminal phosphatase domains. This hypothetical model posits that one signal has cell-wide effects on GLR-1 transport whereas the other has only local effects. Together, these roles could contribute to both homeostatic and Hebbian mechanisms. Overall, this presents and interesting model in which a singular protein modulates both global output and local content of glutamate receptors providing a unified mechanism to control synaptic plasticity and neuronal excitation.

## Material and Methods

### *C. elegans* culture and strains

*C. elegans* strains were kept on NGM and fed the *E. coli* strain OP50 at 20° C. Double and triple mutants were generated by standard genetic methods. Transgenic strains were created by microinjection of *lin-15 (n765ts)* worms with plasmids containing the *lin-15p::lin-15* rescue or in other appropriate genetic backgrounds using pCT61 encoding *egl-20p::nls::DsRed* to express DsRed in the nucleus of four epithelial cells in the tail to visualize rescued worms (Hoerndli et al., 2013). Strains and PCR primers used are all described in the Key Resources table and additional information on the nature of the genetic alleles used are shown in Table 1 below.

### C. elegans transgenes

*akIs201*, *rig-3p::SEP::mCherry::GLR-1; akIs154; rig-3p::HA::glr-1::Dendra2*; *CsfEx14, ptp-3p::PTP-3A::HA; csfEx75, flp-18p::PTP-3A::HA; CsfEx131, flp-18p::PTP-3A::HA; CsfEx132, flp-18p::PTP-3A-Nterm::HA; csfEx134, flp-18p::PTP-3A-Cterm::HA; CsfEx135, hsp16-2p::PTP-3A::HA*.

### Plasmids and cloning

All generated plasmids, except *rig-3p::SEP::mCherry::GLR-1*, *rig-3p::GLR-1::Dendra2* and the original *ptp-3p::PTP-3A::HA*, were made by the Takara In-Fusion cloning method. Plasmids were created by PCR linearization and subsequent insertion of desired sequence by complimentary overhangs created by the PCR linearization. Primers were designed using Takara’s In-Fusion primer design tool and synthesized by Integrated DNA Technology.

The *ptp-3p::PTP-3A* plasmid was generously donated by Dr. Brian Ackley. Using the Takara Bio’s In-Fusion PCR primer tool, the *ptp-3p* promoter was swapped for the AVA specific *flp-18p* promoter. This plasmid was then used to generate the *flp-18p::PTP-3A N-term* keeping the first 232 amino acids and deleting the rest using In-Fusion primers. This stretch of amino acids includes the signal peptide followed by 2 Ig-Like domains and then a STOP. A similar strategy was used for *flp-18p::PTP-3A C-term*, in which In-Fusion primers led to the retention of base pairs 5895-7656 (from the ATG) or a stretch of 556 amino acids. The same replacement strategy using In-Fusion as the one described for *flp-18p* was used for swapping in the *hsp-16p* promoter for inducible heat-shock. All the primers used for cloning are listed in Supplemental Table 1.

### Confocal Imaging

Imaging was conducted on a spinning disk confocal microscope (Olympus IX83) equipped with 488 and 561 nm excitation lasers (Andor ILE Laser Combiner). Images were captured using an Andor iXon Ultra EMCCD camera through a 100x/1.40 oil objective (Olympus). Devices were controlled remotely for image acquisition using MetaMorph 7.10.1 (Molecular Devices).

#### Transport imaging

All imaging was performed on worms containing the *akIs201* array in *glr-1* null (*ky176*) background. Worms were mounted on a 10% agarose pad with 1.6 µl of a 1:1 mixture of polystyrene beads (Polybead, catalog #00876-15, Polysciences) and 30 µM Muscimol (catalog #195336, MP Biomedicals). Using a microcoverslip, worms were positioned such that the AVA neuron was near the coverslip to allow for clear visualization. Through the 100X objective, the neurons were located using the 561 nm excitation laser; then, a cross section in the proximal portion of the AVA neurites was photobleached using a 3 W 488 nm Coherent solid-state laser Coherent solid-state laser (Genesis MX MTM) set to 0.5 W output and a pulse time of 1 s targeted using a Mosaic II digital mirror device (Andor Mosaic 3). After photobleaching, a 500-frame stream was collected at 100 ms exposure per frame with the 561 nm excitation laser in a single Z-plane. Kymographs were generated by the Kymograph tool in MetaMorph with a 20-pixel line width as described in (Doser et al., 2020; Hoerndli et al., 2013).

#### FRAP

Worms containing akIs201 were mounted as described in the “Transport Imaging” section. A proximal region of AVA just distal of the bifurcation of AVAL and AVAR was used as the imaging region and this location was saved for each worm using MetaMorph’s stage position memory function. An image stack was acquired using the 488 nm and 561 nm excitation lasers at 500 ms exposure (20 total images every 0.25 um starting 2.5 µm below to 2.5 µm above the neurites). Around 60 µm to the left and right of the region was photobleached using the same parameters described earlier. Finally, using the stage memory position, the initial process location was photobleached and then immediately imaged with both excitation lasers. Imaging was repeated at 2, 4-, 8-, 12- and 16-minute time points post photobleach.

#### Photoconversion

All photoconversion was performed on worms containing the *akIs154* (GLR-1::Dendra2) array in *glr-1*null (*ky176*) backgrounds. A region containing 2-3 synaptic puncta was converted using an ROI selection tool and a custom control journal to target a 500 ms pulse of 35 mW/mm^2^ from a 405 nm laser (475 mW, LDC8, Power Technologies Inc.) using a Mosaic II digital mirror device (Andor Mosaic 3). Immediately following photoconversion, 20 images were taken using both 561 nm and 488 nm excitation lasers starting 2.5 µm below and finishing 2.5 µm above the process with 500 ms exposure This was repeated at times 2, 4-, 8-, 12-, and 16-minutes post-conversion.

### Short-term and long-term associative olfactory memory

Assays were conducted as described previously (Hadziselimovic et al., 2014). Animals were synchronized using the egg preparation method (Stiernagle, 2006) and then grown on 10 cm NGM plates with OP50. One-day old adult animals were collected, washed three times in CTX buffer, then subjected to different treatment conditions corresponding to either naïve, 1x conditioning, STAM, 2x conditioning or LTAM described below. For all conditions, the chemotaxis of the animals after treatment was tested by placing 50-200 worms in the center of a 10 cm chemotaxis plate (CTX 5 mM KH_2_PO_4_/K_2_HPO_4_ [pH 6.0], 1 mM CaCl_2_, 1 mM MgSO_4_, 2% agar) with 1 µl of 1 M Sodium Azide, and either 0.1% diacetyl (DA) in EtOH or EtOH alone on opposite sides of the plate. After 1 hour at room temperature with closed lids, animals immobilized at the 0.1% Diacetyl, at the EtOH and outside of the spots were counted. The Chemotaxis Index (CI) was calculated as follows: CI = (No. Animals 0.1% DA - No. Animals EtOH) / Total number of animals as described (Bargmann et al., 1993). Counting of animals on the plates was done blind to genotype. The chemotaxis assay was conducted in triplicate for each genotype and condition during a given behavior session and repeated a minimum of 3 sessions. Data from all chemotaxis assays for each condition and genotype were combined to generate the average CI per genotype and condition. For STAM, 50-200 worms were conditioned for 1 hour without food in the presence of 2 µl Diacetyl on the top of the lid of a 10 cm CTX plate. Animals were then either tested immediately (1x cond) or left for 1 hour on NGM with OP50 and tested for STAM (1h memory). For LTAM, the 1-hour conditioning was repeated with a 30-minute interval in between conditioning sessions where worms were able to roam on plates with food. After the second conditioning, worms were either tested immediately for chemotaxis or put on NGM plates with OP50 for 24 hours before LTAM was tested (24h memory; Figure 3A).

### Image analysis

#### Transport

Transport event quantification was performed blinded to the genotype and manually counted using kymographs generated in MetaMorph. Transport velocities and stopping were blindly quantified by manually tracing transport events and subsequently analyzed using the KymoAnalyzer ImageJ plugin (Neumann et al., 2017).

#### FRAP

Image stacks from all time points were converted to maximum projections using MetaMorph’s stack arithmetic function. Average fluorescence from all timepoints were analyzed using the region measurement tool of ImageJ. The ROI chosen included all the puncta clearly seen in the image when left and right peripheral regions were photobleached. The average fluorescence of the same ROI defined before photobleaching was quantified in all subsequent time points. To account for residual fluorescence immediately post photobleach, the average fluorescence of timepoint 0 was subtracted from each subsequent timepoint’s average fluorescence.

#### Dendra

Image stacks from all timepoints were converted to max projections using MetaMorph’s stack arithmetic tool. Photoconverted GLR-1 within a ∼50 μm section of the AVA neurite was quantified using ImageJ’s region measurement tool by outlining the converted region (ImageJ Version: 2.0.0-rc-69/1.53i). The same area measurement was used for all timepoints of the same image. To account for initial red fluorescence of Dendra2 immediately before photoconversion, the average fluorescence of timepoint “before” was subtracted from each subsequent timepoint’s average fluorescence.

### Statistical Analyses

For all analyses with less or equal to three groups a student’s t-test was used. For all microscopy data analyses with more than 3 groups except FRAP and Dendra2 photoconversion, and data with equal variations an ordinary one-way ANOVA with Dunnett’s multiple testing correction was used to determine significance. To determine if the FRAP recovery curve and Dendra2 fluorescence removal were significantly different between genotypes an extra sum-of-squares *F* test applied to the nonlinear hyperbolic regression fit (Figure 4C and 4E) or exponential fit (Figure 4H) to the data. For analyses of short-term and long-term associative memory (STAM and LTAM), a two-way ANOVA with Bonferroni correction for multiple testing was used. All statistics were done use Prism Version 9.1.2 or higher.

## Supporting information

Key Resources Table

## ACKNOWLEDGEMENTS

We would like to thank Dr. Ackley for his generous gifts of the PTP-3A::HA plasmid as well as the College of Veterinary and Biomedical Science at Colorado State University for financial support through an internal College Research Council grant award to Dr. Hoerndli 2020-2021 and NIH NS115947. We would also like to thank Dr. Stacher Hoerndli for the graphic model in Figure 7.

## AUTHOR CONTRIBUTIONS

DP and FJH, conceived and designed all experiments except behavioral associative learning experiments for the manuscript. DP did all experiments and recorded data for the manuscript except for the associative learning behavior. Imaging of synaptic GLR-1 after heat shock was performed and analyzed by DP, KK and ZL. PCR analysis of PTP-3 mRNA in *ptp-3(mu256)* was done by ZL and FJH. Molecular cloning and creation of transgenic strains was conducted by DP with aid from RD. AS designed, recorded, and analyzed the data for all associative learning behavior experiments. FJH wrote the manuscript. AS, DP, RD, KK and ZL gave feedback and input on the manuscript.

## Supplemental Material

### Supplemental Figures

**Supplemental Figure 1:**
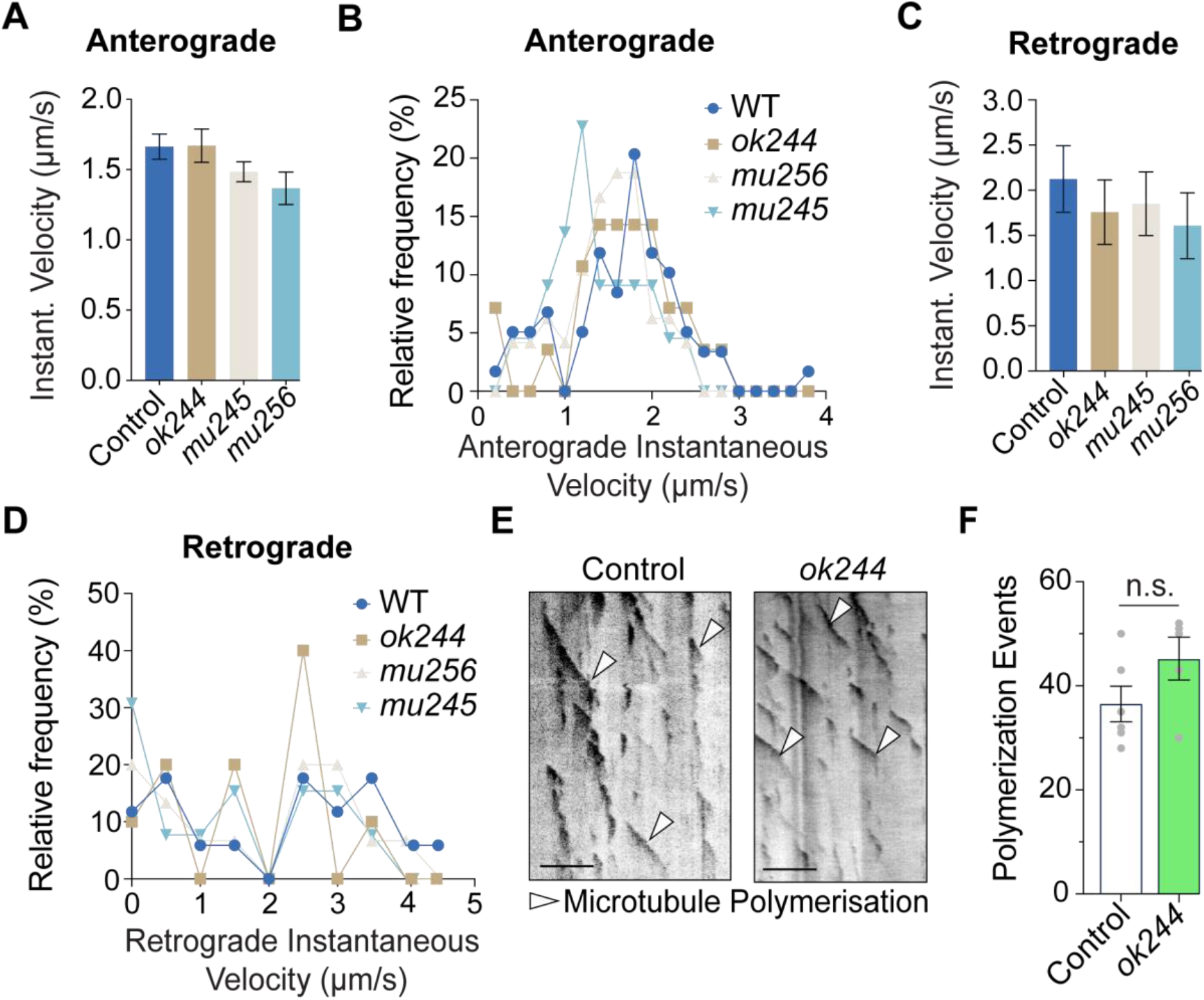
Loss of PTP-3A does not affect the average velocity of GLR-1-containing vesicles or microtubule dynamics. All transport quantifications (A-D) are based on transport of mCherry::GLR-1 from SEP::mCherry::GLR-1 expression in AVA in the *glr-1(ky176)* genetic null background (see methods). A) Average instantaneous anterograde velocities of GLR-1 transport events. B) Distribution of instantaneous anterograde velocities of GLR-1 transport. Control (n=59 transport events), *ok244* (n=28 events), *mu256* (n=48 events) and *mu245* (n=22 events). C) Average instantaneous retrograde velocities of GLR-1 transport events. D) Distribution of instantaneous retrograde velocities of GLR-1 transport. Control (n=17 transport events), *ok244* (n=10 events), *mu256* (n=15 events) and *mu245 (n=13 events)*. E) Representative kymographs of microtubule polymerization and **F)** quantification of events using EBP-2::GFP. Control (n=6), *ok244* (n=5) animals analyzed. An unpaired student’s t-test was used in F. Error bars represent SEM and all scale bars represent 5 µm.

**Supplemental Figure 2:**
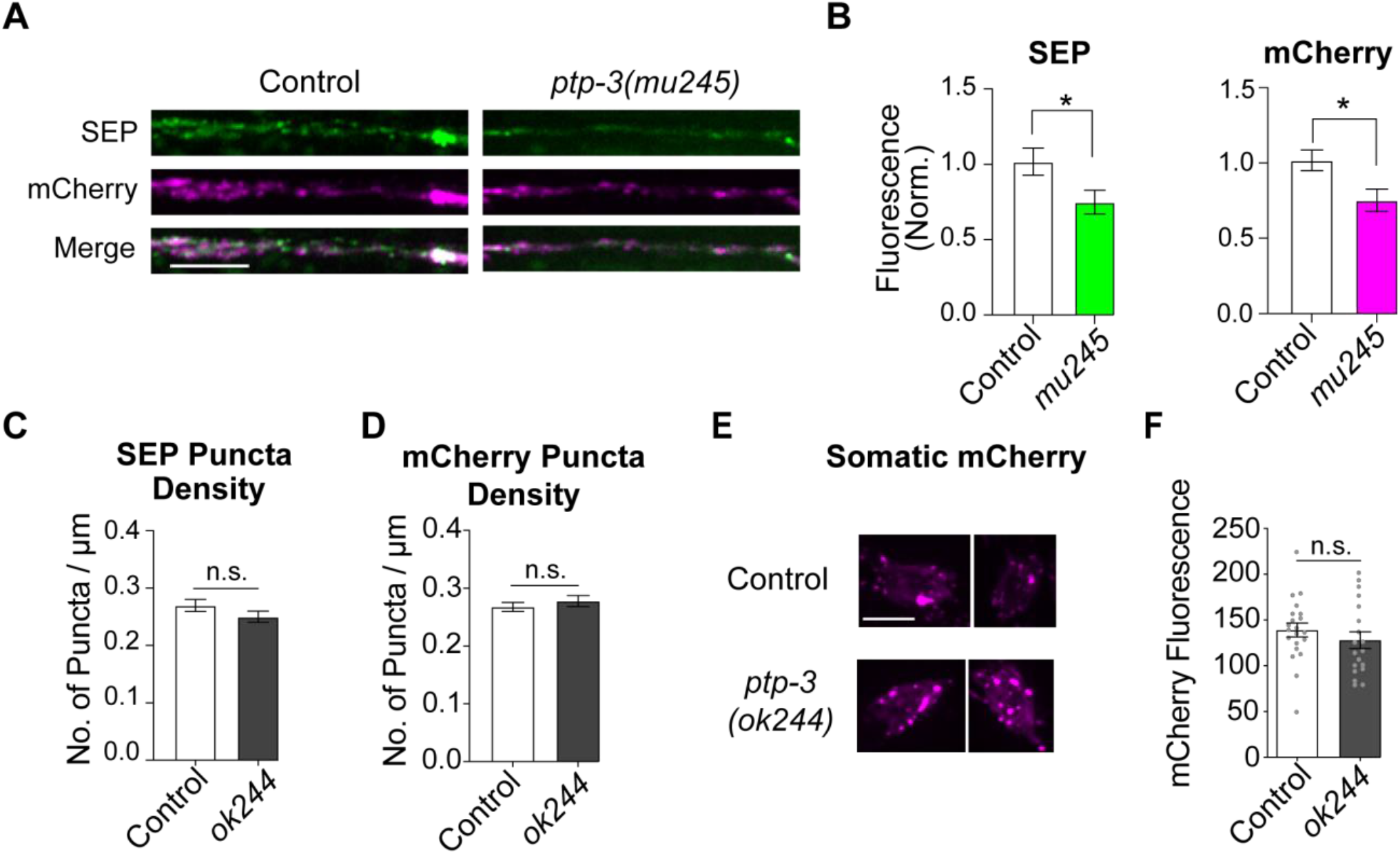
Loss of PTP-3A alters GLR-1 synaptic content but does not affect somatic GLR-1 expression. **A)** Representative confocal images of SEP and mCherry fluorescence in the proximal region of AVA expressing SEP::mCherry::GLR-1 in control and *mu245* animals in the *glr-1(ky176)* null background. **B**) Quantification of synaptic SEP and mCherry fluorescence, normalized to same day control animals. Control (n=54) and *mu245* (n=17). Unpaired student’s t-test. *< p=0.05. **C)** Quantification of SEP and **D)** mCherry puncta number per µm. **E)** Representative confocal maximum projections of image stacks of mCherry fluorescence from SEP::mCherry::GLR-1 in the cell bodies of AVA neurons. **F)** Somatic mCherry fluorescence quantification of AVA GLR-1. Control (n=21) and *ok244* (n=19) cell bodies. Unpaired student’s t-test. Error bars represent SEM and all scale bars represent 5 µm.

**Supplemental Figure 3:**
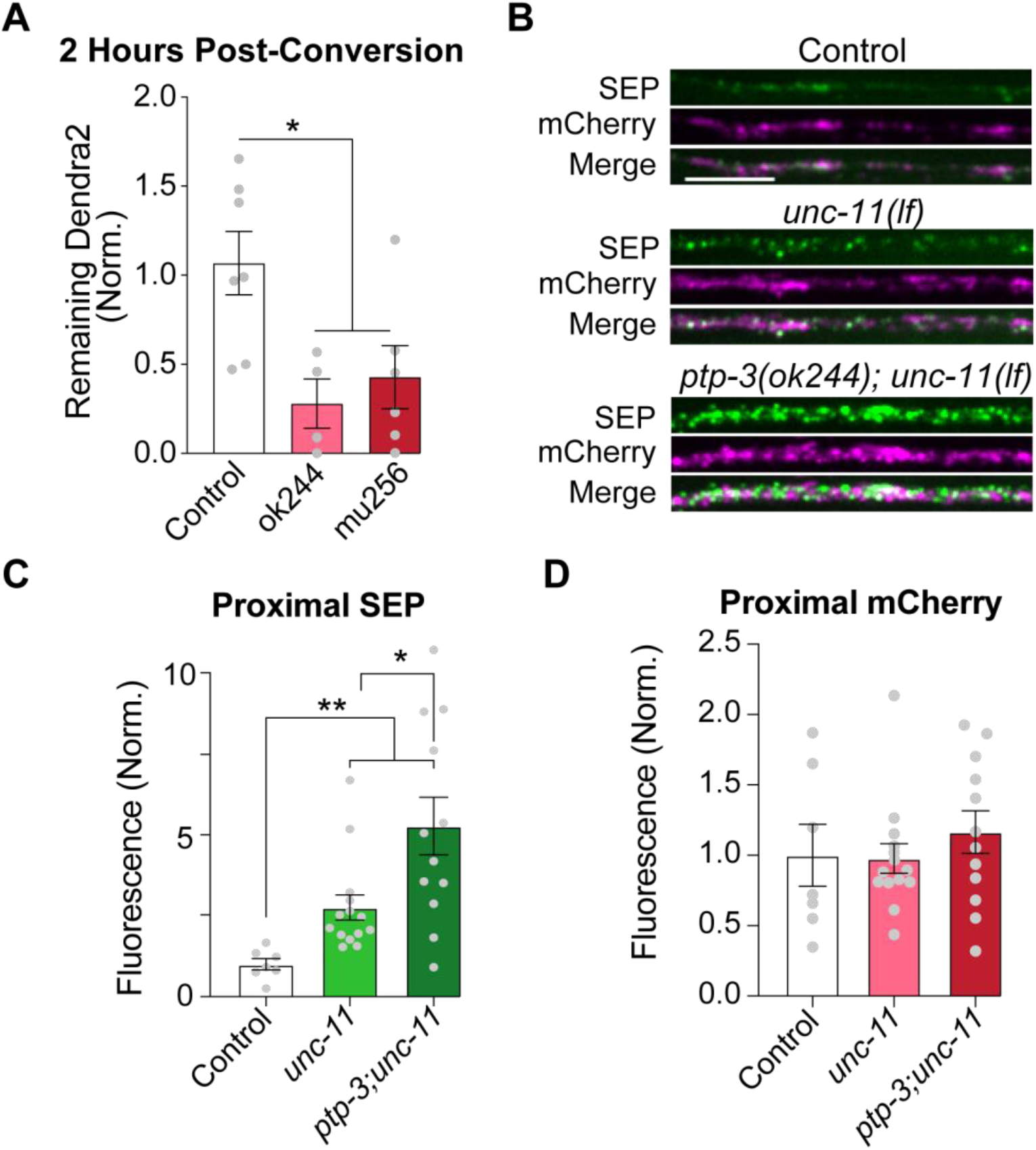
Retention of GLR-1 at synapses depends on PTP-3A and adaptin-dependent internalization. **A)** Percent of original photoconverted GLR-1::Dendra2 signal remaining 2 hours after photoconversion in the proximal region of AVA. Control (n=7), *ptp-3(ok244)* (n=4), and *ptp-3(mu256)* (n=6), * *p* <0.05. **B)** Images of SEP and mCherry fluorescence of SEP::GLR-1::mCherry at proximal AVA synapses in control, *unc-11(lf) and unc-11(lf); ptp-3(ok244)*. **C-D)** Quantification of the SEP and mCherry synaptic fluorescence from images in B. Control (n=7), *unc-11(lf) (n=14), unc-11(lf); ptp-3(ok244)* (n=12). Unpaired Student’s t-test: * *p* < 0.05 and ** *p* < 0.01. Error bars are SEM and scale bars = 5 µm.

**Supplemental Table 1:**
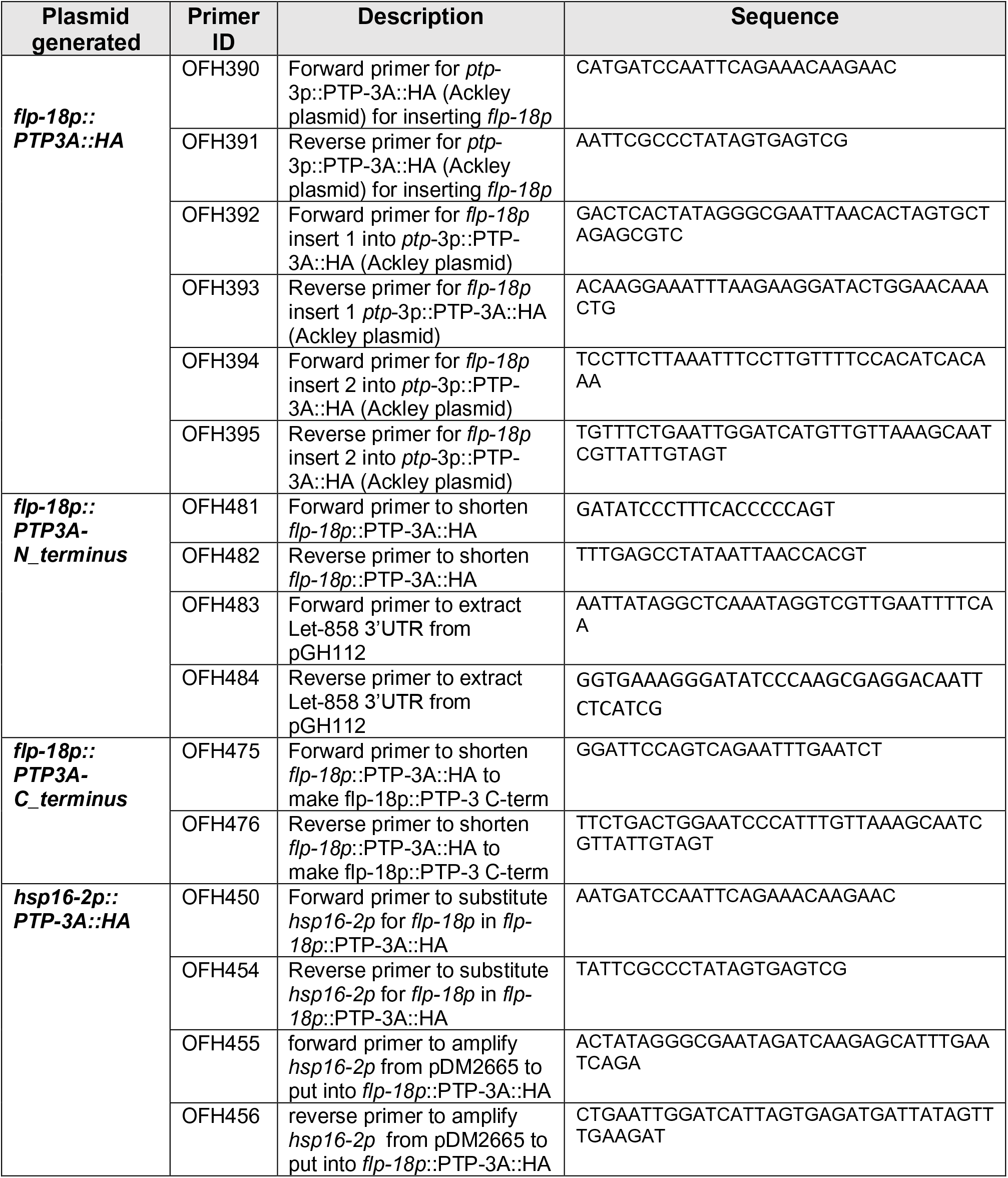
Names, description and sequences of primers used to generate plasmids for *C. elegans* expression.

## REFERENCES

Ackley, B.D., Harrington, R.J., Hudson, M.L., Williams, L., Kenyon, C.J., Chisholm, A.D., and Jin, Y. (2005). The two isoforms of the Caenorhabditis elegans leukocyte-common antigen related receptor tyrosine phosphatase PTP-3 function independently in axon guidance and synapse formation. Journal of Neuroscience 25, 7517–7528.

Aicher, B., Lerch, M.M., Müller, T., Schilling, J., and Ullrich, A. (1997). Cellular redistribution of protein tyrosine phosphatases LAR and PTPσ by inducible proteolytic processing. Journal of Cell Biology 138, 681–696.

Bargmann, C.I., Hartwieg, E., and Horvitz, H.R. (1993). Odorant-selective genes and neurons mediate olfaction in C. elegans. Cell 74, 515–527.

Borcel, E., Palczynska, M., Krzisch, M., Dimitrov, M., Ulrich, G., Toni, N., and Fraering, P.C. (2016). Shedding of neurexin 3β ectodomain by ADAM10 releases a soluble fragment that affects the development of newborn neurons. Scientific Reports 6, 1–12.

Burbea, M., Dreier, L., Dittman, J.S., Grunwald, M.E., and Kaplan, J.M. (2002). Ubiquitin and AP180 regulate the abundance of GLR-1 glutamate receptors at postsynaptic elements in C. elegans. Neuron 35, 107–120.

Callender, J.A., and Newton, A.C. (2017). Conventional protein kinase C in the brain: 40 years later. Neuronal Signaling 1.

Caylor, R.C., Jin, Y., and Ackley, B.D. (2013). The Caenorhabditis elegans voltage-gated calcium channel subunits UNC-2 and UNC-36 and the calcium-dependent kinase UNC-43/CaMKII regulate neuromuscular junction morphology. Neural Dev 8, 10.

Coles, C.H., Jones, E.Y., and Aricescu, A.R. (2015). Extracellular regulation of type IIa receptor protein tyrosine phosphatases: Mechanistic insights from structural analyses. Seminars in Cell and Developmental Biology 37, 98–107.

Dai, Y., Taru, H., Deken, S.L., Grill, B., Ackley, B., Nonet, M.L., and Jin, Y. (2006). SYD-2 Liprin-α organizes presynaptic active zone formation through ELKS. Nature Neuroscience 9, 1479–1487.

Debant, A., Serra-Pagès, C., Seipel, K., O’Brien, S., Tang, M., Park, S.H., and Streuli, M. (1996). The multidomain protein Trio binds the LAR transmembrane tyrosine phosphatase, contains a protein kinase domain, and has separate rac-specific and rho-specific guanine nucleotide exchange factor domains. Proc Natl Acad Sci U S A 93, 5466–5471.

Diering, G.H., and Huganir, R.L. (2018). The AMPA Receptor Code of Synaptic Plasticity. Neuron 100, 314–329.

Doser, R.L., Amberg, G.C., and Hoerndli, F.J. (2020). Reactive Oxygen Species Modulate Activity-Dependent AMPA Receptor Transport in C. elegans. The Journal of Neuroscience 40, JN-RM-0902-20.

Dunah, A.W., Hueske, E., Wyszynski, M., Hoogenraad, C.C., Jaworski, J., Pak, D.T., Simonetta, A., Liu, G., and Sheng, M. (2005). LAR receptor protein tyrosine phosphatases in the development and maintenance of excitatory synapses. Nature Neuroscience 8.

Emperador-Melero, J., de Nola, G., and Kaeser, P.S. (2021). Intact synapse structure and function after combined knockout of ptpδ, ptpσ, and lar. Elife 10, 1–15.

Feinberg, E.H., VanHoven, M.K., Bendesky, A., Wang, G., Fetter, R.D., Shen, K., and Bargmann, C.I. (2008). GFP Reconstitution Across Synaptic Partners (GRASP) Defines Cell Contacts and Synapses in Living Nervous Systems. Neuron 57, 353–363.

Gladding, C.M., Collett, V.J., Jia, Z., Bashir, Z.I., Collingridge, G.L., and Molnár, E. (2009). Tyrosine dephosphorylation regulates AMPAR internalisation in mGluR-LTD. Molecular and Cellular Neuroscience 40, 267–279.

Groc, L., and Choquet, D. (2020). Linking glutamate receptor movements and synapse function. Science 368.

Grochowska, K.M., Andres-Alonso, M., Karpova, A., and Kreutz, M.R. (2022). The needs of a synapse—How local organelles serve synaptic proteostasis. The EMBO Journal 41.

Gutiérrez, Y., López-García, S., Lario, A., Gutiérrez-Eisman, S., Delevoye, C., and Esteban, J.A. (2021). KIF13A drives AMPA receptor synaptic delivery for long-term potentiation via endosomal remodeling. J Cell Biol 220.

Hadziselimovic, N., Vukojevic, V., Peter, F., Milnik, A., Fastenrath, M., Fenyves, B.G., Hieber, P., Demougin, P., Vogler, C., de Quervain, D.J., et al. (2014). Forgetting is regulated via Musashi-mediated translational control of the Arp2/3 complex. Cell 156, 1153–1166.

Han, K.A., Um, J.W., and Ko, J. (2019). Intracellular protein complexes involved in synapse assembly in presynaptic neurons. In Advances in Protein Chemistry and Structural Biology, (Academic Press), pp. 347–373.

Hangen, E., Cordelières, F.P., Petersen, J.D., Choquet, D., and Coussen, F. (2018). Neuronal Activity and Intracellular Calcium Levels Regulate Intracellular Transport of Newly Synthesized AMPAR. Cell Reports 24, 1001–1012.e3.

Hanus, C., and Ehlers, M.D. (2016). Specialization of biosynthetic membrane trafficking for neuronal form and function. Curr Opin Neurobiol 39, 8–16.

Harrington, R.J., Gutch, M.J., Hengartner, M.O., Tonks, N.K., and Chisholm, A.D. (2002a). The C. elegans LAR-like receptor tyrosine phosphatase PTP-3 and the VAB-1 Eph receptor tyrosine kinase have partly redundant functions in morphogenesis. Development 129, 2141–2153.

Hartin, S.N., Hudson, M.L., Yingling, C., and Ackley, B.D. (2015). A synthetic lethal screen identifies a role for lin-44/Wnt in C. elegans embryogenesis. PLoS ONE 10, e0121397.

Hendricks, M., Ha, H., Maffey, N., and Zhang, Y. (2012). Compartmentalized calcium dynamics in a C. elegans interneuron encode head movement. Nature 487, 99–103.

Herring, B.E., and Nicoll, R.A. (2016). Kalirin and Trio proteins serve critical roles in excitatory synaptic transmission and LTP. Proceedings of the National Academy of Sciences 113, 2264–2269.

Hirbec, H., Francis, J.C., Lauri, S.E., Braithwaite, S.P., Coussen, F., Mulle, C., Dev, K.K., Couthino, V., Meyer, G., Isaac, J.T.R., et al. (2003). Rapid and differential regulation of AMPA and kainate receptors at hippocampal mossy fibre synapses by PICK1 and GRIP. Neuron 37, 625–638.

Hoerndli, F.J., Walser, M., Hoier, E.F., de Quervain, D., Papassotiropoulos, A., and Hajnal, A. (2009). A conserved function of C. elegans CASY-1 calsyntenin in associative learning. PLoS ONE 4.

Hoerndli, F.J., Maxfield, D.A., Brockie, P.J., Mellem, J.E., Jensen, E., Wang, R., Madsen, D.M., and Maricq, A. v. (2013). Kinesin-1 regulates synaptic strength by mediating the delivery, removal, and redistribution of AMPA receptors. Neuron 80, 1421–1437.

Hoerndli, F.J., Wang, R., Mellem, J.E., Kallarackal, A., Brockie, P.J., Thacker, C., Madsen, D.M., and Maricq, A. v. (2015). Neuronal activity and CaMKII regulate kinesin-mediated transport of synaptic AMPARs. Neuron 86, 457–474.

Hoogenraad, C.C., Milstein, A.D., Ethell, I.M., Henkemeyer, M., and Sheng, M. (2005). GRIP1 controls dendrite morphogenesis by regulating EphB receptor trafficking. Nat Neurosci 8, 906–915.

Hoogenraad, C.C., Feliu-Mojer, M.I., Spangler, S.A., Milstein, A.D., Dunah, A.W., Hung, A.Y., and Sheng, M. (2007). Liprinα1 Degradation by Calcium/Calmodulin-Dependent Protein Kinase II Regulates LAR Receptor Tyrosine Phosphatase Distribution and Dendrite Development. Developmental Cell 12, 587–602.

Huganir, R.L., and Nicoll, R.A. (2013). AMPARs and Synaptic Plasticity: The Last 25 Years. Neuron 80, 704–717.

Juo, P., Harbaugh, T., Garriga, G., and Kaplan, J.M. (2007). CDK-5 regulates the abundance of GLR-1 glutamate receptors in the ventral cord of Caenorhabditis elegans. Molecular Biology of the Cell 18, 3883–3893.

Kennedy, M.J., Davison, I.G., Robinson, C.G., and Ehlers, M.D. (2010). Syntaxin-4 defines a domain for activity-dependent exocytosis in dendritic spines. Cell 141, 524– 535.

Kim, W.H., Watanabe, H., Lomoio, S., and Tesco, G. (2021). Spatiotemporal processing of neural cell adhesion molecules 1 and 2 by BACE1 in vivo. Journal of Biological Chemistry 296, 100372.

Kobayashi, H., Saito, T., Sato, K., Furusawa, K., Hosokawa, T., Tsutsumi, K., Asada, A., Kamada, S., Ohshima, T., and Hisanaga, S.I. (2014). Phosphorylation of Cyclin-dependent kinase 5 (Cdk5) at Tyr-15 is inhibited by Cdk5 activators and does not contribute to the activation of Cdk5. Journal of Biological Chemistry 289, 19627–19636.

Kolkman, M.J.M., Streijger, F., Linkels, M., Bloemen, M., Heeren, D.J., Hendriks, W.J.A.J., and Van Der Zee, C.E.E.M. (2004). Mice lacking leukocyte common antigen-related (LAR) protein tyrosine phosphatase domains demonstrate spatial learning impairment in the two-trial water maze and hyperactivity in multiple behavioural tests. Behavioural Brain Research.

Lua, W., Isozaki, K., Roche, K.W., and Nicoll, R.A. (2010). Synaptic targeting of AMPA receptors is regulated by a CaMKII site in the first intracellular loop of GluA1. Proc Natl Acad Sci U S A 107, 22266–22271.

Miller, K.E., DeProto, J., Kaufmann, N., Patel, B.N., Duckworth, A., and Van Vactor, D. (2005). Direct observation demonstrates that Liprin-α is required for trafficking of synaptic vesicles. Current Biology 15, 684–689.

Mitchell, C.J., Kim, M.S., Zhong, J., Nirujogi, R.S., Bose, A.K., and Pandey, A. (2016). Unbiased identification of substrates of protein tyrosine phosphatase ptp-3 in C. elegans. Molecular Oncology 10, 910–920.

Padamsey, Z., McGuinness, L., Bardo, S.J., Reinhart, M., Tong, R., Hedegaard, A., Hart, M.L., and Emptage, N.J. (2016). Activity-Dependent Exocytosis of Lysosomes Regulates the Structural Plasticity of Dendritic Spines. Neuron 93, 132–146.

Sclip, A., and Südhof, T.C. (2020). LAR receptor phospho-tyrosine phosphatases regulate NMDA-receptor responses. Elife 9, 1–21.

Serra-Pages, C., Saito, H., and Streuli, M. (1994). Mutational analysis of proprotein processing, subunit association, and shedding of the LAR transmembrane protein tyrosine phosphatase. Journal of Biological Chemistry 269, 23632–23641.

Serra-Pagès, C., Kedersha, N.L., Fazikas, L., Medley, Q., Debant, A., and Streuli, M. (1995). The LAR transmembrane protein tyrosine phosphatase and a coiled-coil LAR-interacting protein co-localize at focal adhesions. The EMBO Journal 14, 2827–2838.

Serra-Pagès, C., Medley, Q.G., Tang, M., Hart, A., and Streuli, M. (1998). Liptins, a family of LAR transmembrane protein-tyrosine phosphatase-interacting proteins. Journal of Biological Chemistry 273, 15611–15620.

Setou, M., Seog, D.-H., Tanaka, Y., Kanai, Y., Takei, Y., Kawagishi, M., and Hirokawa, N. (2002). Glutamate-receptor-interacting protein GRIP1 directly steers kinesin to dendrites. Nature 417, 83–87.

Shah, K., and Rossie, S. (2018). Tale of the Good and the Bad Cdk5: Remodeling of the Actin Cytoskeleton in the Brain. Molecular Neurobiology 55, 3426–3438.

Shin, H., Wyszynski, M., Huh, K.H., Valtschanoff, J.G., Lee, J.R., Ko, J., Streuli, M., Weinberg, R.J., Sheng, M., and Kim, E. (2003). Association of the kinesin motor KIF1A with the multimodular protein liprin-α. Journal of Biological Chemistry 278, 11393–11401.

Stetak, A., Hörndli, F., Maricq, A. V., van den Heuvel, S., and Hajnal, A. (2009). Neuron-Specific Regulation of Associative Learning and Memory by MAGI-1 in C. elegans. PLoS ONE 4, e6019.

Stiernagle, T. (2006). Maintenance of C. elegans. WormBook 1–11.

Streuli, M., Krueger, N.X., Thai, T., Tang, M., and Saito, H. (1990). Distinct functional roles of the two intracellular phosphatase like domains of the receptor-linked protein tyrosine phosphatases LCA and LAR. The EMBO Journal 9, 2399–2407.

Streuli, M., Krueger, N.X., Ariniello, P.D., Tang, M., Munro, J.M., Blattler, W.A., Adler, D.A., Disteche, C.M., and Saito, H. (1992). Expression of the receptor-linked protein tyrosine phosphatase LAR: proteolytic cleavage and shedding of the CAM-like extracellular region. EMBO J 11, 897–907.

Tan, H.L., Chiu, S.L., Zhu, Q., and Huganir, R.L. (2020). GRIP1 regulates synaptic plasticity and learning and memory. Proc Natl Acad Sci U S A 117, 25085–25091.

Trotter, J.H., Hao, J., Maxeiner, S., Tsetsenis, T., Liu, Z., Zhuang, X., and Südhof, T.C. (2019). Synaptic neurexin-1 assembles into dynamically regulated active zone nanoclusters. Journal of Cell Biology 218, 2677–2698.

Uetani, N. (2000). Impaired learning with enhanced hippocampal long-term potentiation in PTPdelta-deficient mice. The EMBO Journal 19, 2775–2785.

Venkatesh, H.S., Johung, T.B., Caretti, V., Noll, A., Tang, Y., Nagaraja, S., Gibson, E.M., Mount, C.W., Polepalli, J., Mitra, S.S., et al. (2015). Neuronal activity promotes glioma growth through neuroligin-3 secretion. Cell 161, 803–816.

Venkatesh, H.S., Tam, L.T., Woo, P.J., Lennon, J., Nagaraja, S., Gillespie, S.M., Ni, J., Duveau, D.Y., Morris, P.J., Zhao, J.J., et al. (2017). Targeting neuronal activity-regulated neuroligin-3 dependency in high-grade glioma. Nature 549, 533–537.

Volk, L., Chiu, S.L., Sharma, K., and Huganir, R.L. (2015). Glutamate Synapses in Human Cognitive Disorders. Annual Review of Neuroscience 38, 127–149.

Vukojevic, V., Gschwind, L., Vogler, C., Demougin, P., de Quervain, D.J., Papassotiropoulos, A., and Stetak, A. (2012). A role for alpha-adducin (ADD-1) in nematode and human memory. EMBO J 31, 1453–1466.

Vukojevic, V., Mastrandreas, P., Arnold, A., Peter, F., Kolassa, I.T., Wilker, S., Elbert, T., de Quervain, D.J.F., Papassotiropoulos, A., and Stetak, A. (2020). Evolutionary conserved role of neural cell adhesion molecule-1 in memory. Translational Psychiatry 10.

Williams, S.L., Lutz, S., Charlie, N.K., Vettel, C., Ailion, M., Coco, C., Tesmer, J.J.G., Jorgensen, E.M., Wieland, T., and Miller, K.G. (2007). Trio’s Rho-specific GEF domain is the missing Gαq effector in C. elegans. Genes and Development 21, 2731–2746.

Won, S.Y., and Kim, and H.M. (2018). Structural Basis for LAR-RPTP–Mediated Synaptogenesis. Molecules and Cells 41, 622–630.

Wyszynski, M., Kim, E., Dunah, A.W., Passafaro, M., Valtschanoff, J.G., Serra-Pagès, C., Streuli, M., Weinberg, R.J., and Sheng, M. (2002). Interaction between GRIP and liprin-α/SYD2 is required for AMPA receptor targeting. Neuron 34, 39–52.

Xie, X., Luo, L., Liang, M., Zhang, W., Zhang, T., Yu, C., and Wei, Z. (2020). Structural basis of liprin-α-promoted LAR-RPTP clustering for modulation of phosphatase activity. Nature Communications 11.

Yong, A.J.H., Tan, H.L., Zhu, Q., Bygrave, A.M., Johnson, R.C., and Huganir, R.L. (2020). Tyrosine phosphorylation of the AMPA receptor subunit GluA2 gates homeostatic synaptic plasticity. Proceedings of the National Academy of Sciences 201918436.

