## Supplementary material for "A Dual Role for LAR-RPTP in Regulating Long-distance Transport and Synaptic Retention of AMPARs, Essential for Long Term Associative Memory": Key Resources Table

**KEY RESOURCES TABLE**

| REAGENT or RESOURCE | SOURCE | IDENTIFIER |
|---|---|---|
| **Chemicals:** | | |
| Muscimol | ThermoFisher | ICN19533610 |
| Sodium azide | ACROS organics | 16628-22-8 |
| Low Melt Agarose for microscopy pads | VWR | J234-100G |
| Polystyrene Beads | Polysciences | 00876-15 |
| Critical Commercial Assays | | |
| Plasmid Miniprep-Classic Kit | ZYMO | 11-308AB |
| Takara In-Fusion Cloning Kit | TakaraBio | NC0476444 |
| Zymoclean Gel DNA Recovery Kit | ZYMO | D4008 |
| QIAquick PCR Purification Kit | Qiagen | 28104 |
| **Experimental Models: Organisms/Strains** | | |
| N2 | CGC | |
| *glr-1(ky176) III; akIs201 V* | Maricq | FJH15 |
| *ptp-3(ok244) II; glr-1(ky176) III; akIs201 V* | Hoerndli | FJH83 |
| *ptp-3(mu245) II; glr-1(ky176) III; akIs201 V* | Hoerndli | FJH348 |
| *ptp-3(mu256) II; glr-1(ky176) III; akIs201 V* | Hoerndli | FJH379 |
| *ptp-3(ok244) II; lin15 (n765ts) X; CsfEx75* [pDP1 (20ng/ul) + pBSKS (5ng/ul) + pJM23 (20ng/ul)] | Hoerndli | FJH323 |
| *ptp-3(ok244)II; CsfEx132* [pDP2(20ng/ul) + pBSKS (5ng/ul) +pJM23 (20ng/ul) + pCT61 (25ng/ul)] | Hoerndli | FJH383 |
| *ptp-3(ok244) II; glr-1(ky176) akIs154 III* | Hoerndli | FJH51 |
| *ptp-3(mu256) II; glr-1(ky176) akIs154 III* | Hoerndli | FJH350 |
| *glr-1(ky176) akIs154 III* | Maricq | FJH30 |
| *ptp-3(ok244) II; glr-1(ky176) III; akIs201 V; CsfEx14 [ptp-3p::PTP-3A::HA* (50ng/ul)+ *pBSKS2* (5ng/ul) + *pCT61* (25 ng/ul) | Hoerndli | FJH106 |
| *ptp-3(ok244) II; glr-1(ky176) III; akIs201 V; CsfEx131* [pDP1 (50ng/ul) + pBSKS (25ng/ul) + pCT61 (25ng/ul)] | Hoerndli | FJH321 |
| *ptp-3(ok244) II; glr-1(ky176) III; akIs201 V; CsfEx132* [pDP2 (30ng/ul) + pBSKS (45ng/ul) + pCT61 (25ng/ul)] | Hoerndli | FJH230 |
| *ptp-3(mu256) II; glr-1(ky176) III; akIs201 V; CsfEx134* [pDP4 (20ng/ul) + pBSKS (55ng/ul) + pCT61 (25ng/ul)] | Hoerndli | FJH245 |
| *ptp-3(ok244) II; glr-1(ky176) III; akIs201 V; CsfEx135* [pDP5(20ng/ul) + pBSKS (55ng/ul) + pCT61 (25ng/ul)] | Hoerndli | FJH354 |
| **Genotyping Oligonucleotides:** | | |
| *ptp-3 wild type* forward primer: ATCGTCCTTATCTGCACCTAG | Hoerndli | OFH21 |
| *ptp-3 wild type* reverse primer: CATTGTATTTCGGTGGCTTCCAGG | Hoerndli | OFH23 |
| *ptp-3(ok244) reverse* primer: GATGCAATCCAGTCACCACATG | Hoerndli | OFH22 |
| *ptp-3(mu256)* forward primer: TCGATACGCGAATGTGGCTG | Hoerndli | OFH145 |
| *ptp-3(mu256)* reverse primer: GAAGGCTCCAGTTCTTCCAATTCC | Hoerndli | OFH146 |
| *ptp-3 (mu245)* forward primer: GATCGTGTCTGTATCACTGG | Hoerndli | OFH159 |
| *ptp-3 (mu245) reverse*: TCAATCGGTTCACCATCG | Hoerndli | OFH160 |

| | | |
|---|---|---|
| **Recombinant DNA:** | | |
| *ptp-3p::PTP-3A::HA* | Brian Ackley | |
| *pflp-18p::PTP-3A::HA* | Hoerndli | pDP1 |
| *pflp-18p::PTP-3A_NTerminal 2Ig* | Hoerndli | pDP2 |
| *pflp-18p::PTP-3A_Cterminal* | Hoerndli | pDP4 |
| *hsp16-2p::PTP-3::HA* | Hoerndli | pDP5 |
| *egl-20p::NLS::dsRed* | Maricq | pCT61 |
| *lin-15p::lin-15* | Maricq | pJM23 |
| **Software and Algorithms:** | | |
| MatLab | MathWorks | R2022a |
| Custom Matlab code for synaptic fluorescence | Hoerndli et al., 2015 | |
| FIJI/ImageJ | NIH | ImageJ 1.53q |
| Kymoanalyzer | Neuman et al., 2017 | |
| Metamorph | Molecular Devices | 10.1.161 |
| Prism | GraphPad | 9.1.2 |
| Other | | |

### RESOURCE AVAILABILITY

#### Lead Contact

#### Materials Availability

Plasmids and Strains generated in this study will be made available upon request

#### Data and Code Availability

This study did not generate any unique datasets or code.
